# A self-limiting mechanotransduction feedback loop ensures robust organ formation

**DOI:** 10.1101/2025.09.25.678607

**Authors:** Yusuke Mori, Paul Robin, Anne Belle Briggs, Daniel S. Levic, Jiacheng Wang, Kira L. Heikes, Michel Bagnat, Edouard Hannezo, Akankshi Munjal

## Abstract

Organ morphogenesis uses mechanotransduction feedback loops to convert forces into gene expression changes that regulate cell mechanics. How these loops integrate with developmental programs to ensure robust outcomes remains unclear. We show Yap mechanotransduction establishes a self-limiting positive feedback loop for semicircular canal formation in zebrafish. Canal development proceeds through bud initiation, extension, and fusion within the otic epithelium. Local swelling of hyaluronan-rich extracellular matrix (ECM) in the bud activates Yap in a spatial pattern. Yap induces its target *ccn1l1*, promoting further ECM expansion to sustain bud extension. This feedback loop confers developmental robustness: graded knockdown of *ccn1l1* reduces extension rate, yet canal formation persists and fails only with strong disruption. Critically, the loop contains its own termination mechanism. During bud fusion, PKA-CREB signaling, activated by an adhesion GPCR, *gpr126,* suppresses *ccn1l1*, ending the loop. These findings reveal how mechanotransduction loops with built-in termination provide developmental control by integrating mechanical forces, transcriptional responses, and morphogenetic outcomes.

## Main

Signaling pathways employ sophisticated feedback mechanisms to control tissue morphogenesis – the process by which tissues remodel into complex and functional organs^1,2^. Such feedback interactions can take the form of temporal switches or self-limiting circuits, for example, the morphogen Nodal induces its own inhibitor, Lefty, to establish early embryonic patterning^3,4^. Beyond biochemical signals, developing tissues also experience and respond to physical forces. Mechanotransduction pathways sense forces such as compression, stretching, or shear, and convert them into transcriptional responses that remodel the physical environment, creating feedback loops between mechanics and gene expression during morphogenesis^5,6^.

Yes-associated protein 1 (Yap), a transcriptional co-activator of the Hippo pathway, is a central mechanotransducer^7^. In response to force, Yap and its co-activator Taz translocate to the nucleus, where they bind to the transcription factor Tead to activate genes that regulate proliferation, differentiation, and extracellular matrix (ECM) production^7^. By linking mechanical inputs to transcriptional programs that reshape tissue architecture, Yap provides a molecular basis for feedback between mechanics and gene expression^8,9^. Yet how such mechanotransduction feedback loops are spatially and temporally organized during development and how they are ultimately terminated to avoid unchecked amplification or premature growth restriction remains poorly understood. Understanding these principles is central to explaining how complex organ shapes emerge from the interplay of signals, forces, and transcriptional programs.

The zebrafish semicircular canals (SCCs) enable addressing these questions because they allow for the simultaneous measurement of transcriptional, biochemical, and mechanical dynamics during morphogenesis^10^. SCCs arise from the otic vesicle (OV) epithelium through three discrete steps: initiation of bud formation at six zones, extension of these buds, and the fusion of adjacent bud pairs that results in the formation of three pillar-shaped structures that delineate the three SCCs. Bud initiation is driven by localized synthesis of a hyaluronan (HA)– and Versican–rich ECM that osmotically swells, pushing the overlying cells^10,11^. How this process, initiated in a small group of cells, achieves the systems-level coordination necessary to ensure the buds extend to the right size and fuse successfully remains unknown. We hypothesized that mechanotransduction could provide this coordinating mechanism by linking the local ECM expansion to transcriptional responses and morphogenetic outcomes.

Our work reveals that SCC morphogenesis is driven by a self-limiting positive feedback loop operating through Yap mechanotransduction. Using a combination of quantitative imaging, theoretical modeling, and a live knock-in reporter of endogenous Yap, we observe that ECM expansion mechanically activates Yap in a spatially patterned manner, with peak activity at the bud border, recruiting these cells into the growing bud. Yap in turn induces *cellular communication network factor 1-like 1* (*ccn1l1,* also known as *cyr61l1*), a matricellular protein that promotes HA and Versican synthesis, ensuing rapid ECM expansion. Progressive loss of the Yap-Ccn1l1-ECM loop led to a dose-dependent reduction in bud extension rates, yet fusion remained robust at lower levels of disruption and was abolished only at the highest dose.

Termination of this loop occurs during bud fusion when *ccn1l1* expression declines. Analysis of mutants in a conserved adhesion GPCR, *gpr126*, previously shown to block bud fusion, revealed that *ccn1l1* expression persists, resulting in uncontrolled growth. Mechanistically, the activation of Gpr126 signaling during bud fusion results in phosphorylation and activation of the transcription factor CREB, which suppresses *ccn1l1* expression. Thus, the morphological outcome of positive feedback–bud fusion–serves as a stop signal that terminates the very process it drives–bud extension.

## Results

### ECM swelling-mediated Yap activation is required for bud extension

The SCC in zebrafish develops through the remodeling of the lumenized single-layered embryonic inner ear epithelium, the basement membrane (BM)-enclosed OV, at six “canal-genesis zones” (Fig. 1a)^10,12^. Morphogenesis begins at 45 hours post fertilization (hpf) with the initiation of buds at antero-lateral, postero-lateral, and anterior zones, where cells extend into the lumen with reduced BM contact^10,13^. By 50 hpf, these buds extend, and a posterior bud begins to form. By 60 hpf, antero-lateral and anterior buds fuse, while postero-lateral and posterior buds extend, and ventral and ventro-lateral budding begins. By 72 hpf, all bud pairs fuse to form the anterior, posterior, and ventral pillars, which demarcate the three semicircular canals. We focused our analysis on the anterior buds, which provide a clear example of all morphogenetic steps: pre-budding (40 hpf), initiation (45-48 hpf), extension (48-55 hpf), and fusion (55 hpf).

**Figure 1:**
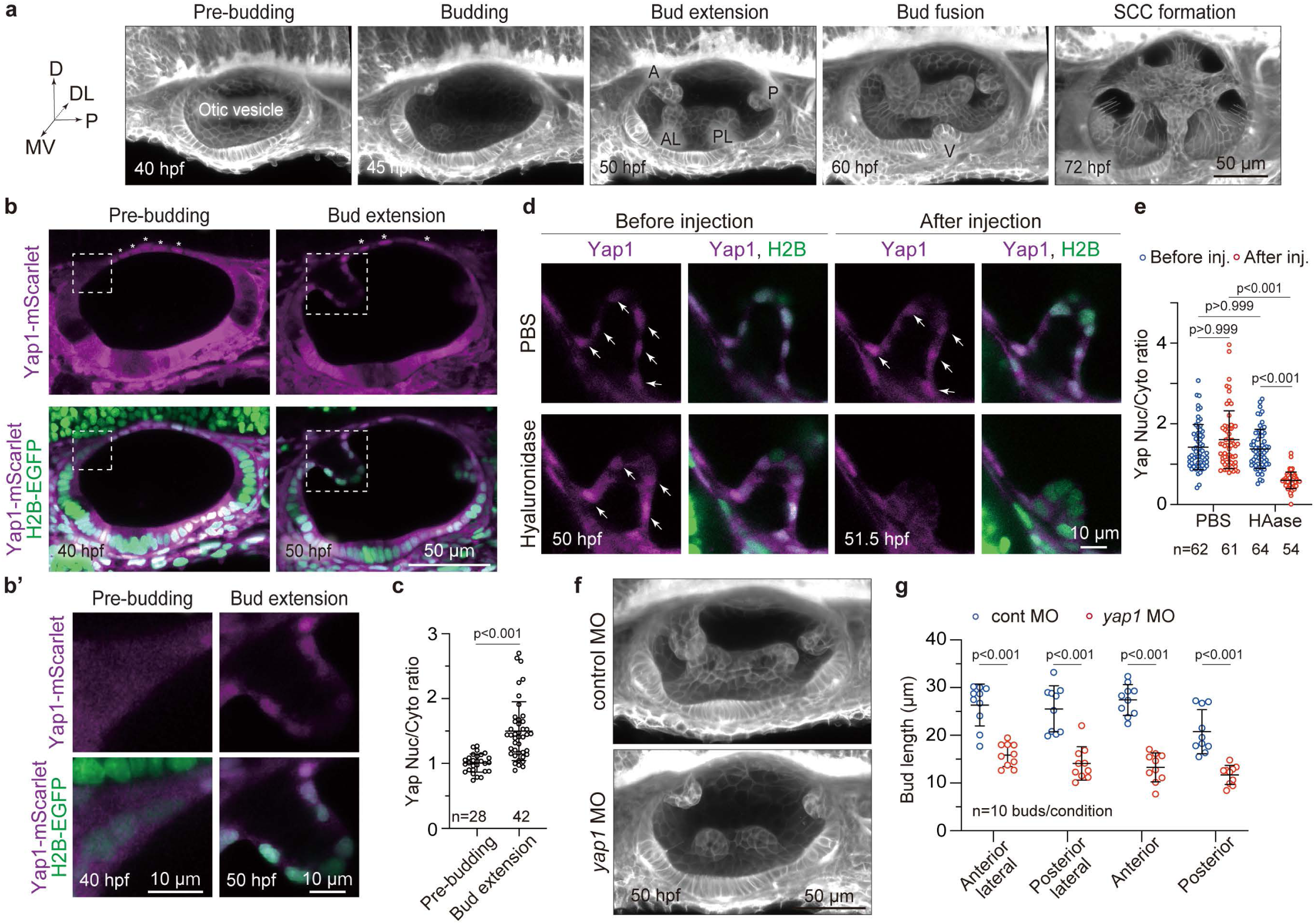
ECM swelling-mediated Yap activation is required for bud extension. (a) Three-dimensional (3D) rendered OVs imaged at selected time points using *Tg(actb2:membrane-neongreen-neongreen)* embryos. A, anterior bud; AL, anterior-lateral bud; P, posterior bud; PL, posterior-lateral bud; V, ventral bud. Developmental stages shown in hours post fertilization (hpf). Scale bar: 50 μm. (b-b’) Two-dimensional (2D) sections of OVs imaged at selected time points using H2B-EGFP mRNA injected *TgKI(Yap1-mScarlet)* embryos. Insets (b’) show enlarged images of the anterior pre-budding or budding region corresponding to the white boxes in (b). Dorsal epithelial cells with nuclear-Yap localization are indicated by asterisks. Scale bar: 10 or 50 μm. Representative images from three independent experiments are shown. (c) Quantification of the Yap nuclear-to-cytoplasmic ratio in the cells at the anterior pre-budding or budding region at selected time points. Data are mean ± SD. *n* denotes the number of cells measured per condition from seven individual embryos from three independent experiments. *P*-value as labeled (Mann-Whitney test). (d-e) Effect of hyaluronidase injection on Yap nuclear translocation. (d) 2D sections of the anterior bud in *TgKI(Yap1-mScarlet)*, injected with H2B-EGFP mRNA, imaged before and at 1.5 hours post-injection (hpi) of PBS or HAase. Pre-injection and post-injection images were acquired from the same individual embryos. Arrows indicate Yap1 translocated to the nucleus. Scale bar: 10 μm. Representative images from two independent experiments are shown. (e) Quantification of the Yap nuclear-to-cytoplasmic ratio in the anterior bud before and at 1.5 hpi of PBS or HAase. Data are mean ± SD. *n* denotes the number of cells measured per condition from ten individual embryos from two independent experiments. *P*-values as labeled (Kruskal–Wallis test with Dunn’s test). (f-g) Effect of *yap1* knockdown on bud growth. (f) 3D-rendered OVs from *Tg(actb2:membrane-neongreen-neongreen)* embryos, injected with control or *yap1* MO, imaged at 50 hpf. Scale bar: 50 μm. Representative images from two independent experiments are shown. (g) Quantification of bud length in embryos injected with control or *yap1* MO at 50 hpf. Data are mean ± SD. *n* denotes the number of buds from individual embryos measured per condition from two independent experiments. *P*-values as labeled (unpaired two-tailed Student’s t-test).

We recently demonstrated that the transcription factor Lmx1b promotes the expression of *hyaluronan synthase 3 (has3)* and *versican b (vcanb),* which encode enzymes for HA and Versican synthesis, respectively, at canal-genesis zones^11^. Charged HA polymers and the proteoglycan Versican that binds HA are secreted basally into the ECM, and osmotic swelling of this HA-rich ECM generates pressure that pushes the overlying epithelium into buds^11^. Longitudinal bud extension is maintained by tangentially distributed actomyosin tethers, known as cytocinches^10^. Because ECM swelling imposes mechanical stress on budding cells^10^ (Extended Fig. 1a-c), we asked whether this stress engages the Yap mechanotransduction pathway.

To characterize the dynamic localization of endogenous Yap protein *in vivo*, we established a *TgKI(Yap1-mScarlet)* reporter line using a knock-in tagging method^14^ (Fig. 1b-b’). Yap activity was quantified by calculating the nuclear-cytoplasmic ratio in budding cells in embryos co-injected with H2B-EGFP mRNA to visualize the nuclei. Prior to budding (40 hpf), Yap was inactive, with a nuclear-to-cytoplasmic ratio around 1. During anterior bud extension at 50 hpf, the ratio increased above 1, indicating active Yap (Fig 1c). We also observed a few cells in the dorsal non-budding region of the OV with a nuclear-to-cytoplasmic ratio greater than 1 at various stages of canal morphogenesis (Fig. 1b and Extended Fig. 1d).

To test whether osmotic swelling of the HA-rich bud-ECM activates Yap in budding cells, we injected hyaluronidase (HAase), an enzyme that degrades HA polymers, into the region surrounding the OV (periotic space). As previously reported, HAase treatment caused shrinkage of the bud-ECM, reduced cell stretching, and inhibited bud extension^10,15^ (Extended Fig. 1e-h). Significantly, nuclear localization of Yap was decreased in HAase-treated embryos, with the nuclear-to-cytoplasmic ratio less than 1 (Fig. 1d-e). HAase treatment had no effect on the nuclear-to-cytoplasmic ratio of non-budding cells in the dorsal OV (Extended Fig. 1i). These findings were further confirmed by Yap immunostaining experiments, in which the nuclear-to-cytoplasmic ratio in budding cells was greater than 1 in controls but reduced to around 1 in HAase-treated embryos (Extended Fig. 1j-m). Together, these results demonstrate that the osmotic swelling of the bud-ECM induces Yap activation in budding cells.

To examine the functional role of Yap, we performed knockdown of *yap1* using a previously characterized morpholino (MO)^16^. Notably, *yap1* knockdown significantly inhibited bud elongation (Fig. 1f-g), resulting in shorter lateral, anterior, and posterior buds at 50 hpf. By 72 hpf, despite the formation of all six buds, 88% of *yap1* morphants failed to initiate pillar formation and consequently did not develop SCCs (Extended Fig. 1n-o). These data demonstrate that Yap is required for successful SCC formation. To uncover the molecular mechanisms underlying this effect, we next examined Yap’s downstream transcriptional targets.

### Yap-target *ccn1l1* expression is Yap-independent at initiation and Yap-dependent during bud extension

To identify downstream targets of Yap, we used available single-cell transcriptomic atlases^17,18^. A zebrafish inner-ear-specific atlas noted expression of a candidate gene, *ccn1l1*, in the SCC-forming cell cluster of the OV, and this expression pattern was validated at the canal-genesis zone during pre-budding and bud extension stages^17,18^. A similar enrichment was observed in the embryo-wide Daniocell atlas^19,20^, where annotation of canal-forming cells using *vcanb* (Versican core protein) revealed *ccn1l1* among the top 10 differentially expressed genes (Fig. 2a–b, Extended Fig. 2a). Ccn1l1 is a paralog of Ccn1 and a member of the CCN family of matricellular proteins^21–23^, which are known downstream targets of Yap^24–27^. Importantly, the 5’ upstream region of *ccn1l1* contains three conserved TEAD-binding motifs recognized by Yap/Taz (GGAATG/CATTCC)^24,28–30^ (Extended Fig. 2b).

**Figure 2:**
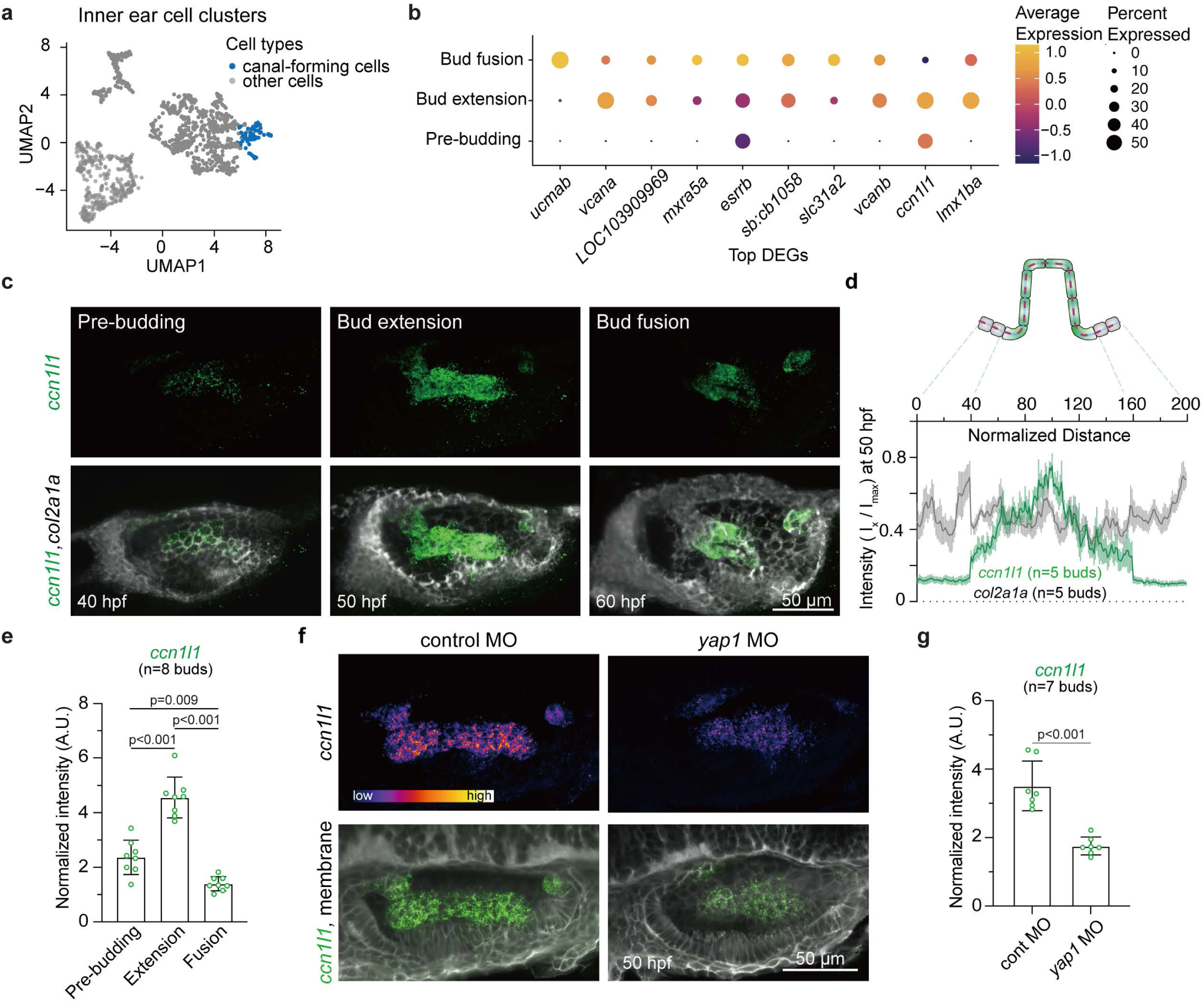
*ccn1l1*, a Yap target gene, is enriched in the canal-genesis zone during SCC formation. (a) UMAP of single-cell RNA of the inner ear cell clusters from the DanioCell dataset. Single cells in the canal-forming cell cluster are colored blue. (b) Dot plot showing the expression levels of the top 10 DEGs in the canal-forming cell cluster during SCC formation: pre-budding (36-40 hpf), bud extension (45-50 hpf), and bud fusion (60-72 hpf) stages. Color encodes the normalized gene expression level, and the dot size encodes the percentage of positive cells within a cluster. (c) 3D-rendered OVs at select time points stained with multiplex *in situ* probes against *ccn1l1* (green) and *col2a1a* (white). Scale bar: 50 μm. Representative images from three independent experiments are shown. (d) Normalized fluorescence probe intensities of *ccn1l1* or *col2a1a* genes across the illustrated region of interest (ROI) at 50 hpf indicated by the red dashed line along the anterior bud. Data are mean ± SEM. *n* denotes the number of buds from individual embryos measured per condition from three independent experiments. (e) Comparison of normalized fluorescence probe intensity of *ccn1l1* in the anterior pre-budding, budding, or pillar region at each stage. Data are mean ± SD. *n* denotes the number of buds or pillars from individual embryos measured from two independent experiments. *P*-values as labeled (one-way ANOVA with Tukey’s test). (f-g) Effect of *yap1* knockdown on *ccn1l1* expression. (f) 3D-rendered OVs from *Tg(actb2:membrane-neongreen-neongreen)* embryos at 50 hpf stained with multiplex *in situ* probes against *ccn1l1* (green). Heatmap represents fluorescence intensity with the same contrast for the *ccn1l1* probe of embryos injected with control and *yap1* MO. Scale bar: 50 μm. Representative images from two independent experiments are shown. (g) Quantification of probe fluorescence intensity of *ccn1l1* probe in anterior buds. Data are mean ± SD. *n* denotes the number of buds from individual embryos measured from two independent experiments. *P*-value as labeled (unpaired two-tailed Student’s t-test).

To quantitatively analyze *ccn1l1* expression dynamics at various developmental stages, we used multiplexed hybridization chain reaction (HCR) fluorescence *in situ* hybridization (FISH)^31^. Consistent with the single-cell atlases^32,18^, *ccn1l1* expression was restricted to the canal-genesis zones (Fig. 2c-d), and was detectable before bud formation at 40 hpf. Its expression increased significantly during bud extension at 50 hpf, and was subsequently downregulated to background levels upon bud fusion at 60 hpf (Fig. 2e). Importantly, MO-mediated *yap1* knockdown led to a 50% reduction in *ccn1l1* expression during bud extension (Fig. 2f-g), making it comparable to levels at the pre-budding stage, confirming that *ccn1l1* is a Yap target. In contrast, *yap1* knockdown did not affect the onset of *ccn1l1* expression at the pre-budding stage (Extended Fig. 2c-d). These findings demonstrate that Yap is essential for increasing *ccn1l1* expression during bud extension, but not for its initial onset.

To understand the initial regulation of *ccn1l1*, we examined its relationship with *lmx1bb*, the transcription factor that, together with its paralog *lmx1ba,* we previously demonstrated patterns HA and Versican synthesis during the pre-budding stage^11^. In the *lmx1bb* null mutants (*lmx1b ^jj410/jj410^*)^31^, which carry a premature stop codon, we observed a 30.1% reduction of *ccn1l1* expression at the pre-budding zone compared to sibling controls (Extended Fig. 2e-f). Pharmacological inhibition of Notch signaling with RO4929097, which expands *lmx1bb* expression^11^, similarly expanded *ccn1l1* expression compared to controls (Extended Fig. 2g).

Altogether, these findings reveal a two-stage regulatory mechanism: Lmx1bb in part initiates *ccn1l1* expression during pre-budding, while Yap amplifies *ccn1l1* expression during extension. Given that Yap sustains bud extension, we hypothesized that *ccn1l1* might promote the synthesis of HA and Versican, which drive bud extension. We next investigated the function of *ccn1l1* to test this possibility.

### Ccn1l1 promotes HA and Versican synthesis to create a positive feedback loop

To investigate the function of *ccn1l1*, we performed CRISPR-Cas9 or MO-mediated *ccn1l1* knockdown. Consistent with its pre-budding expression pattern, the knockdown of *ccn1l1* blocked the onset of budding, resulting in an absence of buds at 50 hpf and no SCCs at 72 hpf (Fig. 3a-b). We next asked how Ccn1l1 drives epithelial budding. CCN proteins regulate cell behaviors such as adhesion, proliferation, and ECM production through their conserved domains^33–35^. Zebrafish Ccn1l1 exhibits high similarity to human CCN1, supporting a conserved function (Extended Fig. 3a-b). We hypothesized that bud formation failure in *ccn1l1* knockdown embryos results from a loss of HA and Versican synthesis. Consistent with this, *has3* and *vcanb* colocalized with *ccn1l1* at canal-genesis zones (Extended Fig. 3c) and exhibited nearly identical temporal expression patterns, with low expression during bud initiation, peaking during bud extension, and returning to baseline during bud fusion, as previously reported^10,11,15^ (Extended Fig. 3d). Indeed, *ccn1l1* knockdown reduced *has3* and *vcanb* expression to background levels at the pre-budding stage and by 50.1% and 73.5%, respectively, at bud extension (Fig. 3c-d and Extended Fig. 3e-f). We conclude that Ccn1l1 drives bud initiation and extension by promoting *has3* and *vcanb* transcription.

**Figure 3:**
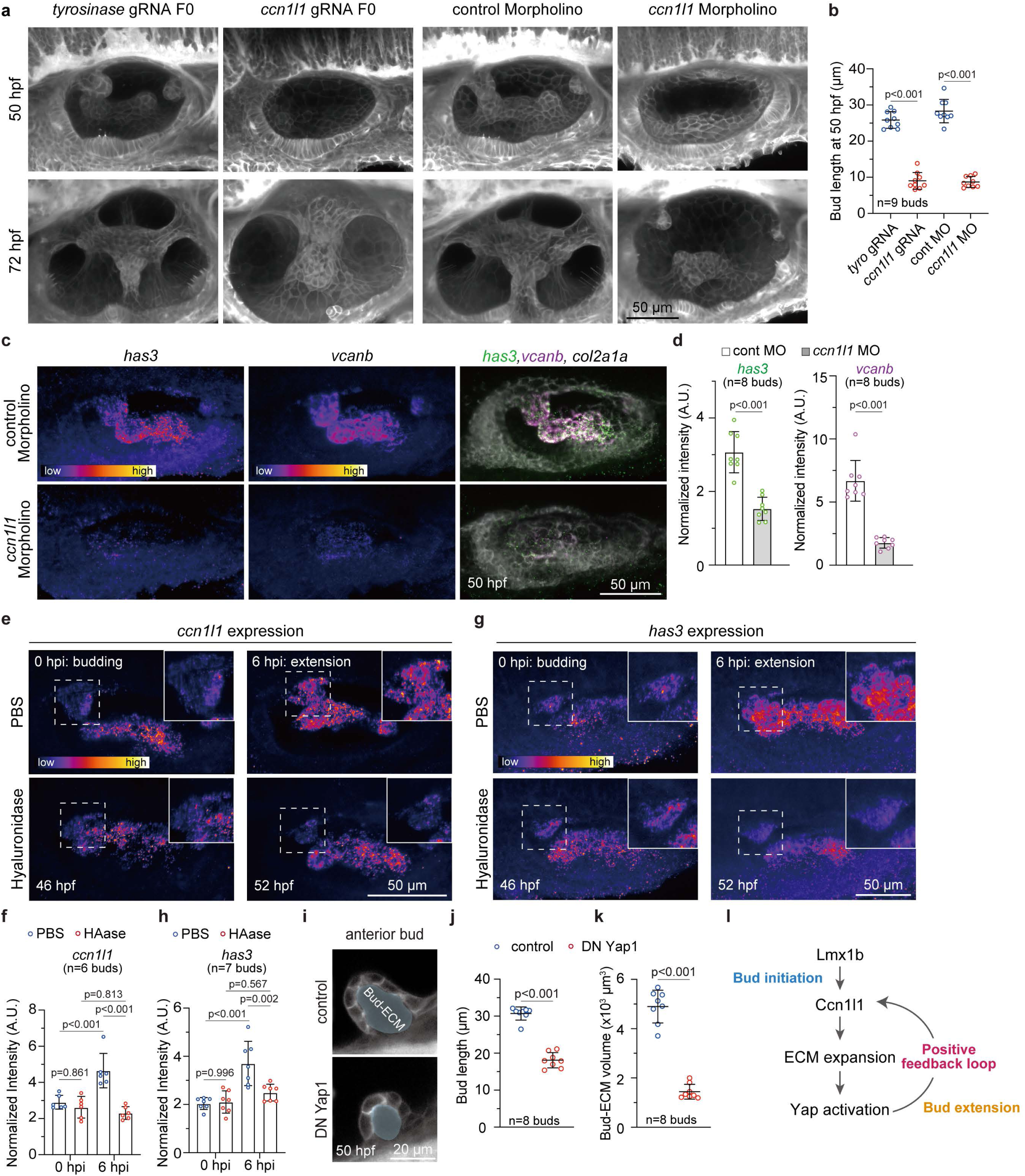
Ccn1l1 induces ECM production and drives the positive feedback loop via Yap mechanotransduction. (a-b) Effect of *ccn1l1* knockdown on bud growth. (a) 3D-rendered OVs from *Tg(actb2:membrane-neongreen-neongreen)* embryos, injected with *tyrosinase* gRNA, *ccn1l1* gRNA, control MO or *ccn1l1* MO, imaged at 50 hpf. Scale bar: 50 μm. Representative images from three independent experiments are shown. *tyrosinase* gRNA is used as a gRNA injection control because it causes loss of pigmentation but does not affect SCC formation. (b) Quantification of bud length in control or *ccn1l1* knockdown embryos at 50 hpf. Data are mean ± SD. In the absence of buds, bud lengths correspond to cell height. *n* denotes the number of buds from individual embryos measured per condition from three independent experiments. *P*-values as labeled (unpaired two-tailed Student’s t-test). (c-d) Effect of *ccn1l1* knockdown on *has3* and *vcanb* expression levels in buds. (c) 3D-rendered OVs at 50 hpf stained with multiplex *in situ* probes against *has3* (green), *vcanb* (magenta), and *col2a1a* (white). Heatmaps represent fluorescence intensity with the same contrast for each probe of embryos injected with control or *ccn1l1* MO. Scale bar: 50 μm. Representative images from two independent experiments are shown. (d) Quantification of probe fluorescence intensity of *has3* or *vcanb* probe in anterior buds. Data are mean ± SD. *n* denotes the number of buds from individual embryos measured per condition from two independent experiments. *P*-values as labeled (unpaired two-tailed Student’s t-test). (e-h) Effect of hyaluronidase injection on *ccn1l1* and *has3* expression levels in buds. 3D-rendered OVs and quantification of probe fluorescence intensity of *ccn1l1* (e-f) or *has3* (g-h) at 0 or 6 hpi of PBS or HAase. Heatmaps represent fluorescence intensity with the same contrast for each probe of embryos injected with PBS or HAase. Scale bar: 50 μm. Representative images from two independent experiments are shown. Data are mean ± SD. *n* denotes the number of buds from individual embryos measured per condition from two independent experiments. *P*-values as labeled (one-way ANOVA with Tukey’s test). (i-k) Effect of DN Yap1 expression on bud extension and bud-ECM volume. (i) 2D sections of anterior buds from *Tg(hsp70l: RFP-DN Yap1)* and sibling control embryos, both *Tg(actb2:membrane-neongreen-neongreen)*, imaged at 50 hpf following heat-shock induction. Regions marked in light blue indicate the bud-ECM region measured in quantifications. Scale bar: 20 μm. Representative images from two independent experiments are shown. Quantification of bud length (j) and bud-ECM volume (k) in sibling control or DN Yap1-expressing embryos at 50 hpf. Data are mean ± SD. *n* denotes the number of buds from individual embryos measured per condition from two independent experiments. *P*-values as labeled (unpaired two-tailed Student’s t-test). (l) A schematic model of the positive mechanotransduction feedback loop during bud growth.

Given the HA-rich bud-ECM’s role in Yap activation and *ccn1l1*’s role in HA production, we hypothesized the existence of a positive feedback loop: *ccn1l1* expression triggers HA production, generating ECM swelling that activates Yap, which in turn enhances *ccn1l1* expression. This predicts that disrupting any component of this loop will affect the others. First, we tested whether disrupting the mechanical force from ECM expansion would affect gene expression. Consistent with our prediction, HAase treatment during bud extension suppressed *ccn1l1* expression to pre-budding levels (Fig. 3e-f) and similarly reduced *has3* expression (Fig. 3g-h). Second, we disrupted transcriptional regulation using a heat-shock inducible dominant-negative (DN) Yap mutant lacking the TEAD-binding domain^36^, preventing the conversion of mechanical cues into transcriptional outputs. These mutants showed significantly reduced bud extension and bud-ECM volume (Fig. 3i-k), confirming that Yap’s transcriptional activity is required for sustained ECM production and morphogenesis.

Together, these complementary approaches, disrupting the mechanical arm with HAase and blocking the transcriptional arm with dominant-negative Yap, demonstrate a positive feedback loop where Ccn1l1 promotes ECM synthesis, ECM expansion activates Yap, and Yap enhances *ccn1l1* expression (Fig. 3l). This creates a self-amplifying system that can drive ECM expansion and bud extension from initially small signals. We next examined the spatiotemporal dynamics of this feedback loop in the growing bud.

### Mechanical patterning of cells at the bud border sustains extension for timely bud fusion

While cell proliferation and convergence-extension through cell rearrangement are typical behaviors underlying tissue growth during development, we previously showed that neither is necessary for bud extension^10^. To determine the source of cells contributing to this bud extension and the role of the Yap-ECM loop, we performed time-lapse imaging and cell tracking. We observed a linear increase in bud cell number over time (Fig. 4a), driven by cells adjacent to the bud that moved toward the initiated bud, turned along the border, and became incorporated into the extending structure (Fig. 4b and Extended Movie 1). Disruption of ECM expansion with HAase treatment or mechanotransduction with *yap1* MO inhibited this increase in cell number (Fig. 4c-d). This suggests that, rather than proliferation, ECM expansion through mechanotransduction exerts pulling forces that recruit adjacent cells into the growing bud.

**Figure 4:**
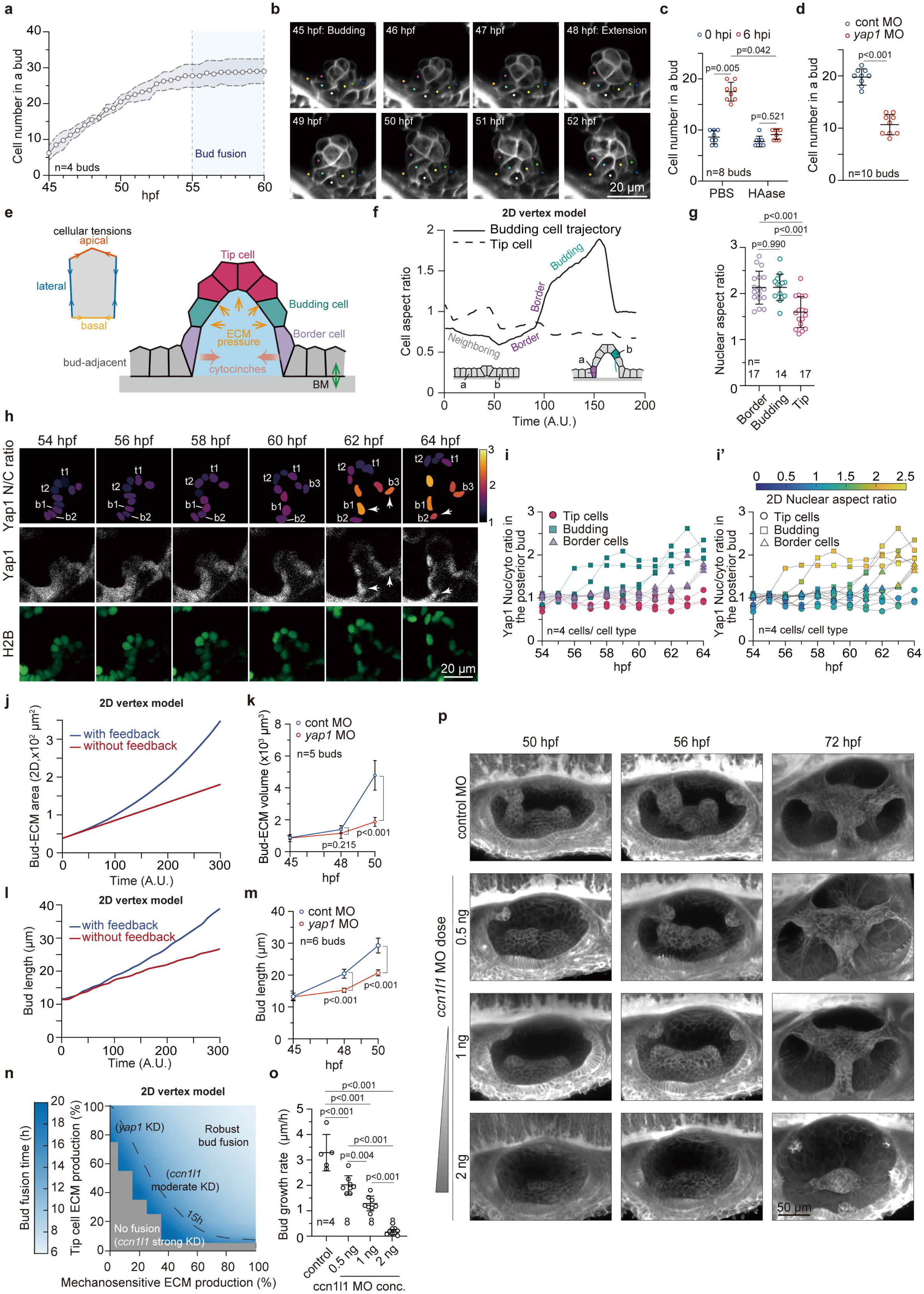
Mechanical patterning by the positive feedback loop recruits adjacent cells to sustain bud extension and ensure timely fusion. (a) Quantification of cell number in the anterior bud across individual time points obtained from time-lapse imaging of *Tg(actb2:membrane-neongreen-neongreen)* embryos. Data are mean ± SD. Dashed lines indicate the range of SD at each time point. *n* denotes the number of buds from individual embryos measured from three independent experiments. (b) 3D-rendered images of an anterior bud of the *Tg(actb2:membrane-neongreen-neongreen)* embryo at selected time points with individual cells tracked. Scale bar: 20 μm. Representative images from two independent experiments are shown. (c) Quantification of cell number in anterior buds of embryos injected with PBS or HAase at 0 or 6 hpi. Data are mean ± SD. *n* denotes the number of buds from individual embryos measured per condition from two independent experiments. *P*-values as labeled (one-way ANOVA with Tukey’s test). (d) Quantification of cell number in anterior buds of embryos injected with control or *yap1* MO at 50 hpf. Data are mean ± SD. *n* denotes the number of buds from individual embryos measured per condition from two independent experiments. *P*-value as labeled (unpaired two-tailed Student’s t-test). (e) Schematics of the 2D vertex model simulations, showing tip, budding and border cells, as well as the relevant physical parameters. All values of physical parameters are given in the supplementary information. (f) Trajectories of the aspect ratio of a tip cell and a recruited cell obtained with 2D vertex model simulations, exhibiting a sharp increase in aspect ratio as cell position shifts from the bud border to the budding region. Illustrations show trajectories of two individual cells from bud initiation to the end of bud extension (defined by the moment where the bud length reaches 30 μm). (g) Quantification of 3D nuclear aspect ratio. The nuclear aspect ratio was calculated as the ratio of the long to short axes (long/short) of the ellipsoidal nucleus. Data are mean ± SD. *n* denotes the number of nuclei from individual cells measured per condition from two independent experiments. *P*-value as labeled (one-way ANOVA with Tukey’s test). (h) Time-lapse analysis of the Yap1 nuclear-to-cytoplasmic (N/C) ratio in a posterior bud of *TgKI(Yap1-mScarlet)*;*Tg(actb2:H2B-EGFP)* embryos at selected time points. Top panels show heatmaps of the Yap1 N/C ratio in individual cells of a posterior bud. Middle and bottom panels show Z-stack images of Yap1-mScarlet and H2B-EGFP, respectively. Nuclear masks used for the Yap1 N/C ratio (top panels) were generated from the H2B-EGFP signal. Yap1 N/C ratios in tip cells (t) and border cells (b) were tracked (top panels). Cells b1, b2, and b3 show sequential Yap activation as they turn the bud border (marked by arrows). Scale bar: 20 μm. Representative images are shown from three independent experiments. (i) Quantification of Yap N/C ratio (i) and 2D nuclear aspect ratio (i’) in tip, budding, and border cells of a posterior bud across individual time points obtained from time-lapse imaging of *TgKI(Yap1-mScarlet)*; *Tg(actb2:H2B-EGFP)* embryos. The N/C ratio of individual cells was calculated every 1 hour and tracked. Heatmap in (i’) represents the 2D nuclear aspect ratio in individual cells. *n* denotes the number of cells from three different posterior buds measured per condition from three independent experiments. (j) Simulation of bud-ECM area over time, with and without mechanotransduction-driven ECM production. Tip cells are set to 100% ECM production rate. (k) Quantification of bud-ECM volume of anterior buds in embryos injected with control or *yap1* MO at selected time points. Data are mean ± SD. *n* denotes the number of cells per condition from two independent experiments. *P*-values at selected time points as labeled (unpaired two-tailed Student’s t-test). (l) Simulation of bud length over time, with and without mechanotransduction-driven ECM production. Tip cells are set to 100% ECM production rate. (m) Quantification of bud length of the anterior buds in embryos injected with control or *yap1* MO at selected time points. Data are mean ± SD. *n* denotes the number of cells per condition from two independent experiments. *P*-values at selected time points as labeled (unpaired two-tailed Student’s t-test). (n) Phase diagram showing the time needed for the bud to reach 30 μm and fuse, as a function of bud ECM production by budding cells (x axis) and tip cells (y axis). The region above the dotted line indicates where fusion happens in less than 15 hours after budding. Grey areas indicate regions where the bud failed to fuse (did not reach 30 μm within 22 hours after budding). (o) Quantification of bud growth rate (μm/hour) of the anterior bud in wild-type embryos or embryos injected with *ccn1l1* MO. Data are mean ± SD. *n* denotes the number of cells per condition from one experiment. *P*-values as labeled (one-way ANOVA with Tukey’s test). Growth rates for each bud were calculated over the following time ranges during bud extension stage: control (48-50 hpf); *ccn1l1* MO 0.5, 1, and 2 ng (50-56 hpf). (p) 3D-rendered OVs from *Tg(actb2:membrane-neongreen-neongreen)* embryos, injected with control or *ccn1l1* MO, imaged at selected time points. Scale bar: 50 μm. Representative images from one experiment per condition are shown.

Because recruitment occurs specifically at the bud border, we hypothesized that mechanotransduction cues are spatially patterned rather than uniform. To explore this, we developed a theoretical vertex model of a 2D epithelial section incorporating experimentally established parameters^10^: apical, lateral, and basal cellular tensions; BM adhesion at the bud border; HA-rich bud-ECM pressure; and anisotropic cytocinch tension orthogonal to the bud’s long axis (Fig. 4e and Extended Fig. 4a-d; see Supplementary Note). The model defines four mechanically distinct regions: bud-adjacent cells (BM-attached), bud border cells (partially BM-attached), budding cells (newly incorporated, BM-detached), and tip cells (leading-edge, BM-detached) (Fig. 4e).

The model indicated that cell recruitment was accompanied by a transient increase in cell aspect ratio (Fig. 4f), with stronger overshoots in border and budding cells compared to tip cells. This spatial pattern was lost when BM attachment or cytocinches were removed (Extended Fig. 4a-d), suggesting that mechanical stress is concentrated at the border cells attached to the BM. To test this prediction, we measured nuclear aspect ratios as markers of cell deformation^37,38^ and found that border and budding cells were more deformed than tip cells (Fig. 4g and Extended Fig. 4e). Long-term time-lapse imaging of the Yap reporter with a nuclear marker confirmed model predictions: cells experienced significant deformation as they flowed toward and turned the bud border, concomitant with strong Yap activation before recruitment (Fig. 4h-i’, Extended Fig. 4f-g, and Extended Movie 2). Moreover, Yap activation occurred sequentially, with each newly arriving adjacent cell activating Yap as it turned the bud border (Fig. 4h and Extended Movie 2). In contrast, nuclear deformation and Yap activity remained low in tip cells (Fig. 4g-i’ and Extended Fig. 4f-g). Notably, *ccn1l1* expression was slightly lower in border cells than in budding cells, consistent with recent Yap activation (Fig. 2d), and its expression domain expanded over time as additional cells were recruited into the extending bud (Extended Fig. 4h-i). Together, these results demonstrate that Yap activitation is spatially restricted to the bud borders and temporally dynamic, with border cells activating Yap and *ccn1l1* expression as they turn the bud corner and are recruited into the bud.

To test the functional significance of spatially patterned mechanotransduction, we modeled bud growth dynamics in which ECM production comprised two components: a constant baseline rate from pre-patterned tip cells and a mechanotransductive contribution proportional to the number of recruited cells. When mechanotransduction strength is high, the model predicts accelerated, non-linear ECM volume growth (Fig. 4j). In contrast, disabling the mechanotransductive contribution, resulted in a slower, more linear increase in ECM volume consistent with constant ECM production from tip cells alone (Fig. 4j). Because bud extension in the model depends on ECM volume growth, reduced ECM production without mechanotransductive contribution led to slower extension rates (Fig. 4l). As predicted by the model, control embryos exhibited rapid ECM growth and faster extension compared to *yap1* MO-injected embryos that showed slower ECM growth and reduced bud extension (Fig. 4k and 4m).

Extending buds eventually meet and fuse to form pillars, completing otic vesicle remodeling and canal demarcation. We next examined how varying extension rates, set by ECM produced by tip cells and by the mechanotransductive recruitment of neighboring cells, determine whether buds achieve fusion. To address this, we generated a phase diagram from ∼500 simulations varying both parameters, with fusion defined as buds reaching 30 µm in length (Fig. 4n), which corresponds to the average distance required for opposing buds to make contact across the otic vesicle lumen in wild-type embryos at 60 hpf (Extended Fig. 5a). The simulations revealed a trade-off: when mechanotransduction strength was low, fusion was delayed and required high tip cell contribution, whereas with high mechanotransduction, even minimal tip cell production was sufficient for bud fusion (Fig. 4n and Extended Fig. 4j). To test these predictions experimentally, we performed dose-dependent MO-knockdowns of *ccn1l1* (Fig. 4o-p and Extended Fig. 4k), which affects both tip cell and mechanotransductive ECM production. At lower doses, extension slowed significantly, but buds still fused successfully by 72 hpf (Fig. 4o-p). At the highest dose, extension dropped below ∼95% of normal rates (Fig. 4o) and buds failed to fuse (Fig. 4p). Thus, mechanotransduction ensures that buds can still achieve fusion across a wide range of tip cell output, with failure occurring only under the most severe loss of ECM production.

Our theoretical model and experimental data demonstrate that mechanical deformation at bud borders activates Yap to drive cell recruitment and ECM expansion, sustaining bud extension rates that ensure successful fusion even when tip cell ECM production varies.

### CREB activation during fusion terminates the positive feedback loop

Intriguingly, *ccn1l1* expression dropped sharply once opposing buds fused (Fig. 2c and 2e). Consistent with this downregulation, both cell recruitment and bud extension plateaued when bud pairs made contact (Fig. 4a and Extended Fig. 5a), indicating a self-termination mechanism that stops further bud growth. A likely candidate for this stop signal is Gpr126 (Adgrg6), a mechanosensitive adhesion-type G-protein-coupled receptor (aGPCR) essential for bud fusion^15,39,40^ and expressed in budding cells throughout morphogenesis^15^ (Extended Fig. 5b). Consistent with previous findings^15^, in *gpr126^st49/st49^* mutants, which carry a premature stop codon^41^, buds fail to fuse and undergo excessive elongation, accompanied by persistent *has3* expression (Fig. 5a-c and Extended Movie 3). We found that *ccn1l1* expression remained persistently elevated in these mutants at both 60 hpf and 72 hpf, contrasting with the normal sharp downregulation upon fusion (Fig. 5b-c and Extended Fig. 5c). This persistent expression drove continued growth: mutant buds were 78.7% longer and contained 76.5% more cells than wild type at 60 hpf (Fig. 5d-e). Notably, bud-ECM volume continued to grow unchecked, with a 464.6% increase in volume (Fig. 5f and Extended Fig. 5d), consistent with the ongoing recruitment of HA-producing cells. This behavior is inconsistent with a genetically predetermined bud cell number, which would have produced a plateau in extension. Instead, this provides further evidence that mechanotransduction recruits new cells to the bud. The 2D vertex model recapitulated this behavior: when fusion was blocked and the mechanotransductive contribution proportional to recruited cells allowed to persist, ECM volume expanded unchecked (Extended Fig. 5e). These results demonstrate that bud fusion provides the essential termination signal for the positive feedback loop; in its absence, mechanotransductive recruitment drives runaway growth.

**Figure 5:**
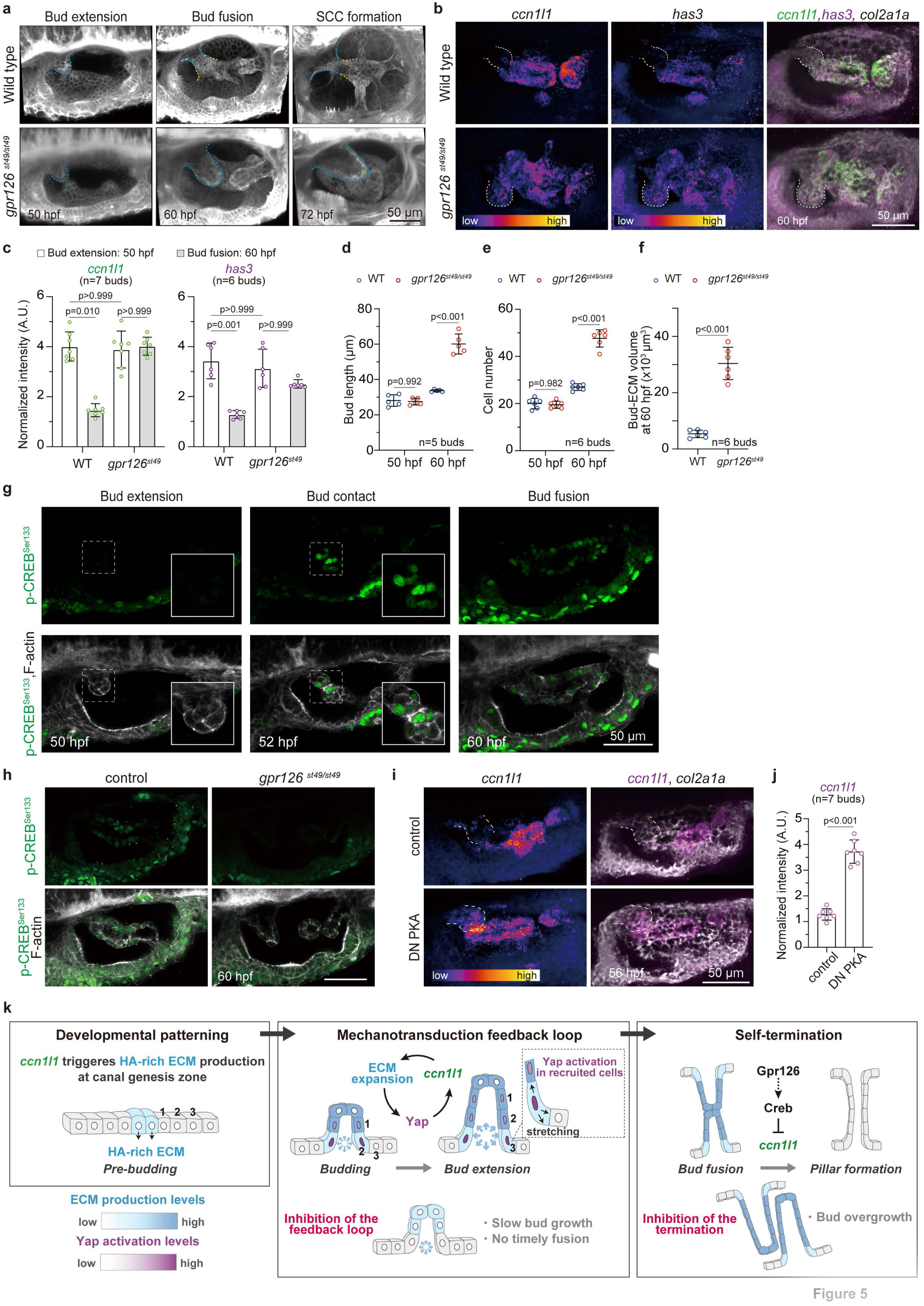
Gpr126-cAMP-PKA-CREB signaling during bud fusion suppresses *ccn1l1* expression to terminate bud extension. (a) 3D-rendered OVs from wild-type or *gpr126 ^st49/st49^* mutant embryos, both *Tg(actb2:membrane-neongreen-neongreen),* imaged at selected time points. Blue and yellow dashed lines indicate the anterior bud/pillar or the anterior-lateral portion of the pillar, respectively. Scale bar: 50 μm. Representative images from two independent experiments are shown. (b-c) *ccn1l1* and *has3* expression levels in *gpr126 ^st49/st49^* mutant embryos. (b) 3D-rendered OVs of wild-type or *gpr126 ^st49/st49^* mutant embryos at 60 hpf stained with multiplex *in situ* probes against *ccn1l1* (green), *has3* (magenta), and *col2a1a* (white). Heatmaps represent fluorescence intensity with the same contrast for each probe of embryos. Scale bar: 50 μm. Representative images from three independent experiments are shown. (c) Quantification of probe fluorescence intensities of *ccn1l1* or *has3* probes in anterior buds or pillars at 50 hpf or 60 hpf. Data are mean ± SD. *n* denotes the number of buds or pillars from individual embryos measured from three independent experiments. *P*-values as labeled (Kruskal-Wallis test with Dunn’s test). (d-f) Effect of *gpr126 ^st49/st49^* mutation on bud growth. Quantification of bud length (d), cell number in the anterior bud or pillar (e), and bud-ECM volume (f) at selected time points. Data are mean ± SD. *n* denotes the number of buds or pillars from individual embryos measured from two independent experiments. *P*-values as labeled (unpaired two-tailed Student’s t-test). (g) 2D sections of OVs showing phospho-CREB and F-actin, stained using anti-phospho-CREB^ser133^ antibody and phalloidin, respectively, at selected time points. Insets show enlarged images of the anterior buds corresponding to the dashed white boxes. Scale bar: 50 μm. Representative images from three independent experiments are shown. (h) 2D sections of OVs showing phospho-CREB and F-actin, stained using anti-phospho-CREB^ser133^ antibody and phalloidin, respectively, at 60 hpf in sibling control or *gpr126 ^st49/st49^* mutant embryos. Scale bar: 50 μm. Representative images from two independent experiments are shown. (i-j) Effect of DN PKA on *ccn1l1* expression in buds. (i) 3D-rendered OVs from *Tg(hsp70l:DN PKA-GFP)* embryos and sibling controls after heat-shock induction, stained with multiplex *in situ* probes against *ccn1l1* (magenta) and *col2a1a* (white). Heatmaps represent fluorescence intensity with the same contrast for each *ccn1l1* probe. Scale bar: 50 μm. Representative images from three independent experiments are shown. (j) Quantification of probe fluorescence intensity of *ccn1l1* in anterior buds or pillars at 56 hpf. Data are mean ± SD. *n* denotes the number of buds or pillars from individual embryos measured from three independent experiments. *P*-value as labeled (unpaired two-tailed Student’s t-test). (k) A schematic model of the Yap mechanotransduction feedback loop and its termination mechanism during semicircular canal formation. HA synthesis is initiated in a small group of cells during the pre-budding stage. As the ECM swells, mechanical stress at the bud border triggers Yap activation, initiating the Yap–*ccn1l1*–ECM positive feedback loop. This loop amplifies the domain by recruiting adjacent cells into the bud, sustaining rapid bud extension and ensuring timely fusion. Disrupting any component of the loop blocks extension. During fusion, Gpr126-PKA-Creb signaling is activated to terminate the feedback loop. Loss of this termination mechanism results in continued ECM production and bud overgrowth.

We next investigated how Gpr126 terminates *ccn1l1* expression. Gpr126 activates the cyclic adenosine monophosphate (cAMP) pathway^42,43^. Pharmacological cAMP elevation with forskolin^44^ strongly reduced both *ccn1l1* and *has3* expression (Extended Fig. 5f-g) and inhibited bud elongation (Extended Fig. 5h-i). Since cAMP signals through Protein Kinase A (PKA), which leads to phosphorylation and activation of the transcription factor CREB^45^, we examined CREB activation during bud fusion. Immunostaining revealed striking CREB phosphorylation at bud-bud contact sites, spreading into budding cells that form the pillar (Fig. 5g). This CREB phosphorylation was absent in *gpr126* mutants (Fig. 5h), demonstrating that Gpr126 signaling mediates the phosphorylation of CREB.

CREB has been reported to downregulate CCN gene transcription in cultured cells^46,47^. To confirm the role of CREB in the inhibition of *ccn1l1*, we expressed DN-PKA lacking the cAMP-binding site^48^. This attenuated CREB phosphorylation and prevented both bud fusion and *ccn1l1* downregulation (Fig. 5i-j and Extended Fig. 5j-k), demonstrating that PKA-CREB signaling is necessary for feedback termination.

Together, these results demonstrate that the Gpr126-cAMP-PKA-CREB pathway suppresses *ccn1l1* and terminates ECM synthesis during bud fusion. This creates a morphological transition-triggered, self-limiting mechanism where the successful completion of morphogenesis (fusion) terminates the positive feedback loop (Fig. 5k).

## Discussion

Our findings reveal how robust SCC morphogenesis emerges from a self-limiting positive feedback mechanism where morphological success triggers its own termination (Fig. 5k). We demonstrate that the mechanotransducer Yap, activated by a HA-rich ECM, drives the expression of the matricellular protein Ccn1l1, which further promotes HA and Versican synthesis to sustain ECM expansion and bud extension. This positive feedback loop is spatially restricted, with Yap activity high at the bud border. Critically, the Gpr126-cAMP-PKA-CREB pathway is activated during bud fusion and terminates *ccn1l1* expression, preventing runaway growth. Thus, the completion of morphogenesis serves as the signal to terminate the very process that drives it.

Most studies of Yap/Taz mechanotransduction have emphasized their role in negative feedback loops that maintain tissue homeostasis, where mechanical forces activate Yap/Taz to promote behaviors that ultimately dissipate those same forces^28,49–51^. More recently, however, it has become clear that Yap/Taz can also participate in positive feedback loops^52^ to amplify mechanical cues rather than inhibiting them. Here we show that Yap activation promotes ECM synthesis, which generates mechanical forces that further sustain Yap activtity to generate self-reinforcing growth dynamics. We propose that this positive feedback mechanism may be important during developmental windows where weak initial signals must be robustly amplified to drive rapid morphogenetic outcomes.

The identification of Ccn1l1 as a key mediator linking Yap activation to ECM production represents a significant advance in understanding how CCN proteins contribute to mechanotransduction networks. While CCN family proteins have been implicated in developmental processes, wound healing, and inflammation^21–23^, their specific role in creating positive feedback loops between mechanical forces and matrix synthesis has not been previously demonstrated. The two-stage regulation of Ccn1l1—initial Lmx1b-dependent expression followed by Yap-dependent amplification—suggests an elegant mechanism for transitioning from developmental patterning to mechanically-driven growth amplification. Notably, *ccn1l1* is also expressed in other organs, including somite and notochord^21^, raising the possibility that similar feedback-driven mechanisms operate more broadly during vertebrate development.

CCN proteins are known to interact with integrins to regulate ECM production^33–35^, and among them, β1 integrin is a well-documented receptor for CCN family members^26,34,53^. β1 integrin expression has indeed been reported in the OV of both zebrafish and mice^54,55^, suggesting that *ccn1l1* may act via integrin to promote ECM production during SCC formation. Beyond ECM regulation, CCN proteins can also remodel the actomyosin cytoskeleton through integrin-Rho GTPase signaling, thereby promoting cellular protrusions and generating mechanical tension^26^. We previously reported that cytocinches progressively increase during bud extension and are essential for longitudinal growth^10^, coinciding with the rise in *ccn1l1* expression. These observations raise the possibility that *ccn1l1* not only promotes ECM expansion, but may also contribute to bud extension by modulating cytoskeletal tension.

Our theoretical model predicts that BM attachment is crucial for spatially restricting Yap activation to border cells, yet this prediction remains to be investigated. Integrins, particularly those that bind laminin components of the BM^56^, are prime candidates for transducing mechanical forces at these sites. Focal adhesion kinase (FAK) and related signaling molecules, such as vinculin, likely serve as mechanosensitive intermediates that link integrin engagement to Yap activation^7^. The precise mechanical threshold required for Yap activation, and how BM composition and stiffness modulate this threshold, represents an important avenue for future investigation. Understanding these molecular details could reveal how tissues fine-tune their mechanosensitivity during morphogenesis.

Adhesion GPCRs (aGPCRs) perform diverse developmental roles in many organs like heart, sciatic nerve, cartilage, and vascular systems, and their dysfunction can lead to organ malformation or embryonic lethality^39,57–60^. Previous work implicated Gpr126 in downregulating ECM production during SCC formation^15^, but the molecular mechanism was unknown. We find that Gpr126 signaling is activated during fusion and terminates ECM production by downregulating *ccn1l1*. Such termination mechanisms may represent a conserved principle by which aGPCRs couple tissue remodeling to signaling and transcriptional outcomes during morphogenesis. Although the precise mode of Gpr126 activation remains unclear, it likely involves ligand binding and/or mechanical forces that expose a tethered agonist (Stachel sequence) to trigger downstream signaling^61–64^.

The integration of mechanotransduction feedback loops to morphological outcomes provides a framework for understanding how complex organ shapes emerge through self-organizing principles. Overall, we demonstrate how robust SCC morphogenesis emerges from an interplay of two evolutionarily conserved pathways. The mechanotransduction feedback loop, Yap-Ccn1l1-HA, drives bud extension, and the Gpr126-cAMP-PKA-CREB pathway couples bud fusion to extension termination. This principle likely applies broadly to developmental systems where patterned growth-promoting signals must first be amplified and then precisely shut down.

## Materials and Methods

### Animals

Zebrafish (*Danio rerio*) EK and AB wild-type strains were used in this study. Adult fish were kept on a 14-hour light/10-hour dark cycle. Embryos were collected by crossing female and male adults (3-18 months old) and raised in 28.5 °C egg water. In this study, the following transgenic and mutant lines were used: *Tg(actb2:membrane-neongreen-neongreen)*^10^*, lmx1b ^jj410/jj410^*^38^, *Tg(hsp70I: RFP-DN-Yap1)* (this study), *Tg(hsp70I: DN-PKA-GFP)* (this study), *gpr126^st49/st49^*^41^, *TgKI(Yap1-mScarlet)^pd^*^1280^ (this study), *Tg(actb2: H2B-EGFP)* (this study), and *Tg(βActin: utrophin-mCherry)*^65^. *gpr126^st49/st49^* mutant fish were maintained as an AB strain background, whereas all other transgenic and mutant lines were maintained as an EK strain background.

### Morpholino injection

Standard control MOs and MOs targeting *ccn1l1* (5′-CTCAGAGGAGACATGGTGTTATCCA-3′, translational blocking), *yap1* (5′-AGCAACATTAACAACTCACTTTAGG-3′, splice-modifying between intron 2 and exon 2) ^16^, and *p53* (5′-GCGCCATTGCTTTGCAAGAATTG-3′, translational blocking)^66^ were purchased from Gene Tools. The *ccn1l1* MO target sequence was designed based on the Danio rerio *ccn1l1* sequence (ENSDART00000051336), retrieved from the Ensembl database, using the Gene Tools Oligo design service. For the *ccn1l1* knockdown, embryos at the single-cell stage were injected with 2 ng of *ccn1l1* MO or control MO, together with 1 ng of *p53* MO. For the *yap1* knockdown, 5 ng of *yap1* MO was injected with 2.5 ng of *p53* MO into embryos at the single-cell stage. MO-injected embryos were screened for inner ear phenotype at 40-50 hpf and used for confocal live imaging, HCR-FISH, or immunostaining at 40-72 hpf. To test the dose-dependent effect of *ccn1l1* MO, embryos at the single-cell stage were injected with either standard control MO (2 ng) or *ccn1l1* MO (0.5, 1, or 2 ng), together with *p53* MO. Injected embryos were screened for inner ear phenotype at 50 hpf, followed by confocal live imaging at 50, 56, and 72 hpf. *p53* MO was co-injected with control, *ccn1l1* or *yap1* MO to reduce MO-injection induced apoptosis and non-specific toxity.

### Genome editing by CRISPR-Cas9

#### *ccn1l1* CRISPANT embryo

Single-guide RNAs (sgRNAs) were synthesized as previously reported^11^. Briefly, sgRNAs targeting multiple coding regions of *ccn1l1* were synthesized using MEGAscript SP6 Transcription Kit (Invitrogen, AM1330) and purified with RNA Clean & Concentrator kit (Zymo Research, R1018). For the knockdown of *ccn1l1*, three sgRNAs were pooled together and injected into embryos as previously described^11^. Briefly, embryos at the single-cell stage were injected with 2 nL of injection solution containing pooled sgRNAs (200 ng/μL: 66.6 ng/μL of each sgRNA) and Cas9 protein (PNA Bio, CP-03, 500 ng/μL). F0 crispant embryos were screened for inner ear phenotype at 50-72 hpf and then used for HCR-FISH or confocal live imaging. The following target sites were used to generate sgRNAs: *ccn1l1* sgRNA1: 5’-CGGCGGGCACAGCGCCAGTCGGG-3’; *ccn1l1* sgRNA2: 5’-GTGTGTCCTCAGGTGCAGGGAGG-3’; *ccn1l1* sgRNA3: 5’-AGGTGGCTGGAGCGATAGGGCGG-3’

#### *tyrosinase* CRISPANT embryo

A sgRNA targeting the *tyrosinase* gene^67^ was synthesized as described above. sgRNA (200 ng/μL) and Cas9 protein (500 ng/μL) were injected into embryos at the one-cell stage as described above. The *tyrosinase* crispant embryos were screened for pigmentation loss at 50-72 hpf and used for HCR-FISH or confocal live imaging. *tyrosinase* crispant embryos were used as an sgRNA injection control, resulting in loss of pigmentation without affecting SCC formation^11^. The following target site was used to generate the sgRNA: 5′-GGACTGGAGGACTTCTGGGG-3′.

#### Generation of *TgKI(Yap1-mScarlet)* line

The *TgKI(Yap1-mScarlet)^pd1280^* line was generated using a previously reported knock-in tagging method^14^. As the targeting cassette, a gene fragment spanning from the middle of intron 7 to the last codon of exon 8 was cloned into pUC19-TgKI-MCS-mScarlet-MCS-polyA (Addgene plasmid #174023) using In-Fusion seamless cloning (Takara Bio) with the following primers: *yap1* Fw: 5′-TTCGCTAGCCCGCGGCTATCAGTGCTTTACAGATCATTGTC-3′; *yap1* Rev: 5′-CACCTCGAGTCTAGATAGCCAGGTTAGAAAGTTCTCC-3′. The sgRNA target sites for genomic editing were mutated in the donor plasmid using Q5 site-directed mutagenesis (New England Biolabs, NEB) with the following primers: *yap1* SDM Fw: 5′-ACATCCTCAACGACATGGAGTCAGTGCTGGC-3′; *yap1* SDM Rev: 5′-CGGAGCTCAAAGCTTCCTGCAGGCTGGGCA-3′. A sgRNA targeting *yap1* (5′-GTTGAGGATGTCGGAGCTTAGGG-3′) was synthesized using the HiScribe T7 Quick High Yield RNA Synthesis Kit (NEB) and purified with the Monarch RNA Cleanup Kit (NEB). The injection mix contained PCR donor (20 nM), gRNA (4 μM), Cas9 protein (3 μM; PNA Bio CP-02), and DMSO (1%). The genomic integration site was confirmed by sequencing at the F2 generation, and the line was designated allele number pd1280.

### Construct the generation and injection of plasmid and mRNA

#### *actb2:H2B-EGFP* plasmid

H2B-EGFP was amplified by PCR using Phusion Flash High-Fidelity PCR Master Mix (Thermo Fisher Scientific, F548L) and appropriate primers from a construct harboring its coding sequence. pMTB *actb2:membrane-neongreen-neongreen* construct^10^, which contains the SP6 promoter for mRNA synthesis, was used as the plasmid backbone. The plasmid backbone was digested with BamHI and AgeI. Then, the amplified H2B-EGFP fragment was cloned downstream of the *actb2* promoter and the SP6 promoter in the pMTB backbone using In-Fusion Snap Assembly Master Mix (Takara, 638948). The H2B-EGFP mRNA was synthesized using the mMESSAGE mMACHINE SP6 transcription kit (Invitrogen, AM1340) from the linearized *actb2:H2B-EGFP* plasmid. 2 nL of mRNA (100 ng/μL) was injected into single-cell-stage embryos from *TgKI(Yap1-mScarlet)* parents. Live embryos were observed at 40-56 hpf by confocal microscopy. To generate the *Tg(actb2:H2B-EGFP)* line, 100 pg of Tol2 mRNA and 50 pg of *actb2:H2B-EGFP* plasmid were injected into embryos at the one-cell stage. The following primers were used for PCR: Fw: 5’-CTTGTTCTTTTTGCAGGATCCCGCCACCATGCCAGAGCCAGCGA-3’; Rev: 5’-TTATCATGTCTGGATCACCGGTTTACTTGTACAGCTCGTCCATGC-3’.

#### *hsp70l: RFP-dominant negative Yap1* plasmid

*hsp70l: RFP-DN-Yap1*^36^ construct harboring the I-SceⅠ site for the transgene was a gift from Kenneth Poss (Morgridge Institute, University of Wisconsin-Madison, USA) and was used as a template for cloning. RFP-DN-Yap1 was amplified by PCR using appropriate primers. The plasmid backbone, which contains a heat-shock-inducible promoter (hsp70l) and Tol2 sites, was digested with BamH1 and AgeI and then gel-purified. The amplified DNA fragment was cloned downstream of the *hsp70l* promoter in the plasmid backbone using the In-Fusion system. To generate the *Tg(hsp70l: RFP-DN-Yap1)* line, 100 pg of Tol2 mRNA and 50 pg of *hsp70l: RFP-DN-Yap1* plasmid were injected into embryos at the one-cell stage. The following primers were used for PCR: Fw: 5’-TTGCAGGATCCGCCACCATGGACAACACCGAGGACG-3’; Rev: 5’ TGGATCACCGGTTCAGATACTACTGTGGGCTGGAGGG-3’

#### *hsp70l: dominant negative PKA-GFP* plasmid

M7 pdnPKA-GFP was a gift from Randall Moon (Addgene plasmid # 16716), used as a template for cloning. The cloning was performed, as described above. Briefly, the amplified dnPKA-GFP fragment was cloned downstream of the *hsp70l* promoter in the plasmid backbone. To generate the *Tg(hsp70l: DN-PKA-GFP)* line, 100 pg of Tol2 mRNA and 50 pg of *hsp70l: DN-PKA-GFP* plasmid were injected into embryos at the one-cell stage. The following primers were used for PCR: Fw: 5’-TACAGTTCAGCCATCCTAGGATTTAGGTGACACTATAGAATACAAGCTACT-3’; Rev: 5’-GTTCCGGATACGTATGAATTCTTATTTGTATAGTTCATCCATGCCATGT-3’

### Drug treatment

For hyaluronidase treatment, we injected hyaluronidase (HAase, Sigma Aldrich, H1136) into the periotic space of dechorionated embryos, with slight modifications to a previously reported method^10^. Briefly, HAase (500 units/mL) was injected into embryos twice at a 3-hour interval from 46 hpf, followed by HCR-FISH or confocal live imaging. For live imaging of *TgKI(Yap1-mScarlet)* embryos, HAase was injected into embryos at 50 hpf, followed by imaging at 1.5 hours post-injection. For immunostaining of Yap1 antibody, HAase was injected into embryos at 50 hpf, and embryos were fixed with 4% PFA in PBS at 3 hours post-injection, followed by immunostaining. For RO4929097 (ApexBio, A4005) treatment, dechorionated embryos at 24 hpf were soaked in egg water containing 50 μM inhibitor for 26 hours, followed by HCR-FISH. For forskolin (Cayman Chemical, 1101810) treatment, dechorionated embryos were soaked in egg water containing 50 μM of the drug at 46 hpf for 6 hours, followed by HCR-FISH or confocal live imaging.

### Multiplex *in situ* hybridization chain reaction (HCR)

HCR probes for each gene were obtained from IDT (Oligo Pools, oPools). Hybridization buffer, amplification buffer, wash buffer, and amplifiers were purchased from Molecular Instruments. mRNA sequences used for HCR probe design were retrieved from the NCBI database. HCR probe sets for each mRNA were designed using a home-made Python script^31^. 40 probe oligonucleotides targeting each mRNA were generated and pooled oligonucleotides were used for HCR-FISH. HCR probe sequences are listed in Supplementary Table 1. HCR-FISH experiments were performed as previously described^10,11^. Dechorionated embryos at various stages were fixed with 4% paraformaldehyde (PFA) in PBS and permeabilized with prechilled acetone. The embryos were then incubated with the hybridization buffer at 37°C for 30 min, followed by incubation with the hybridization solution containing probes overnight at 37°C. After probe hybridization, the embryos were incubated overnight at room temperature in an amplification buffer with fluorescent HCR amplifiers (Alexa-488, Alexa-546, or Alexa-647). Embryos were washed four times with 0.1% Tween-20 in 5× SSC buffer (5x SSCT) and stored in 1x PBS at 4°C until confocal imaging. For nuclear staining, HCR-FISH stained embryos were incubated with Hoechst 33342 (Thermo Fisher Scientific, 62249; 1:2000) at 4°C overnight, followed by washing with 1x PBS. Control and experimental samples were processed within the same tube to ensure uniform treatment. This included the same incubation times, reagent concentrations, and handling procedures, ensuring that any differences in fluorescence intensity were due to variations in target abundance rather than experimental variability.

### Immunohistochemistry

Immunohistochemistry was performed as previously described^11^. Briefly, for phospho-CREB staining, the dechorionated embryos were fixed with 4% PFA/PBS for 2 hours at room temperature, followed by permeabilization with pre-chilled acetone at −20 °C for 7 minutes. Embryos were then blocked with 5% bovine serum albumin (BSA; Sigma-Aldrich, A3311) in 1× PBS containing 0.1% Tween-20 (PBT) at room temperature for 1 h, followed by incubation in primary antibody solution containing rabbit anti-phospho-CREB (Ser133) antibody (Cell Signaling Technology, 9198; 1:100) and mouse anti-GFP antibody (Invitrogen, A11120; 1:500) at 4°C overnight. Embryos were washed with 1x PBT for 5 min, thrice, then incubated with a secondary antibody solution containing fluorescently labeled anti-rabbit and anti-mouse antibodies (Thermo Fisher Scientific, A11008, A11035, A21244, and A11001; 1:200) and phalloidin (Thermo Fisher Scientific, A22287; 1:100) at 4°C overnight. Embryos were washed with 1x PBT for 5 min, thrice, and stored at 4°C. For Yap1 staining, embryos fixed with 4% PFA/PBS were permeabilized with 2% Triton X-100/PBS for 2 hours. Embryos were then blocked with 2% BSA in 1× PBS containing 1% DMSO and 0.1% Triton X-100 (PBDT) for 2 hours at room temperature, followed by incubation in a primary antibody solution containing rabbit anti-Yap1 (Proteintech, 13584-1-AP; 1:200) at 4 °C overnight. Embryos were washed with 1× PBDT for 15 min, thrice, then incubated with a secondary antibody solution containing fluorescently labeled anti-rabbit antibody (1:200) and Hoechst 33342 (1:2000) at 4°C overnight. Following this, embryos were washed with 1× PBDT for 15 minutes, thrice.

### Confocal imaging

Confocal imaging was performed with a Zeiss LSM 980 confocal microscope using a C-Apochromat 40×1.2 NA objective for all fluorescence acquisitions as previously described^11^. Briefly, dechorionated live embryos were anesthetized in 1x tricaine and then mounted dorsolaterally using a canyon mount cast in 1.5% agarose dissolved in egg water and covered with a coverslip. Images of *TgKI(Yap1-mScarlet)* embryos were acquired using Airyscan SR mode, followed by image processing using the Zen software (Zeiss). For imaging embryos stained with HCR-FISH or immunohistochemistry, embryos were mounted on 1.5% agarose dissolved in 1x PBS and then subjected to confocal imaging. Image acquisition settings, such as laser power, gain, and exposure, were kept constant and within the sub-saturation range during microscopy to allow for accurate and quantitative sample comparisons. Time-lapse imaging was performed as previously described^10^. Briefly, embryos were immobilized by injecting 500 μM α-bungarotoxin protein in PBS (Thermo Fisher Scientific, B35450) into the heart 30 min before imaging. The embryos were maintained in a custom-built incubator at 28.5°C during time-lapse imaging. Z-stack images were acquired at 30-minute intervals. For time-lapse analysis of Yap1 nuclear translocation, posterior buds of *TgKI(Yap-mScarlet); Tg(actb2:H2B-EGFP)* embryos were used. Z-stack Images were acquired from 54 hpf at 1 hour intervals with Airyscan Multiplex mode (CO-8Y) for fast scanning, which enabled high-speed imaging with enhanced spatial resolution while reducing excitation dose to minimize photobleaching. Acquired images were processed using Zen software. FluoRender, an open-source software^68^, was used for 3D renderings of images.

### Screening and genotyping of mutants

*gpr126 ^st49/49^* mutants, derived from *gpr126 ^st49/+^* parents, were screened based on inner ear phenotypes at 60 hpf^15^ for immunohistochemistry. For Fig. 5a–f and Extended Fig. 5c-d, EK wild-type and *gpr126 ^st49/49^* mutants were obtained from EK wild-type and *gpr126 ^st49/49^* parents, respectively, and subsequently subjected to HCR-FISH and confocal live imaging at 50-72 hpf. To minimize developmental differences between mutant and wild-type embryos, male and female fish were maintained in a mating tank separated by a removable divider overnight, and fertilized eggs were collected within 15 minutes after the divider was removed the following morning. *lmx1bb ^jj410/+^* mutants were genotyped using the derived cleaved amplified polymorphic sequence (dCAPS) method, as previously described^11^. The *lmx1bb ^jj410/jj410^* mutant obtained from *lmx1bb ^jj410/+^* parents was screened based on inner ear phenotypes at 50 hpf, followed by HCR-FISH.

### Heat-shock activation of hsp70l promoter

For experiments using DN-Yap1 mutants, *Tg(Hsp70l: RFP-DN-Yap1)* fish were crossed with *Tg(actb2:membrane-neongreen-neongreen)* fish. Embryos obtained from this cross were heat-shocked at 37°C for 30 min twice at 5-hour intervals at 40 hpf, followed by confocal live imaging at 50 hpf. DN-Yap1 mutants were identified by RFP fluorescence at 5 hours post-heat shock. RFP-negative siblings were used as controls for confocal imaging. For experiments using DN-PKA mutants, *Tg(Hsp70l: DN-PKA-GFP)* fish were crossed with wild-type or *Tg(βActin:utrophin-mCherry)* fish. To induce acute activation of the heat-shock promoter before bud fusion, embryos were heat-shocked at 42°C for 10 min at 48 hpf, followed by HCR-FISH or confocal live imaging at 56 hpf. DN-PKA mutants were identified by GFP fluorescence at 3 hours post-heat shock. GFP-negative siblings were used as controls for HCR-FISH or confocal live imaging. Heat-shock treatment at 42°C for 10 min was confirmed not to affect zebrafish embryonic development noticeably.

### Modeling

See Theory Supplementary Note

### Analysis of single-cell RNA sequencing data

Single-cell RNA sequencing analysis was performed using Daniocell2023_SeuratV4.rds^19,20^ available at https://daniocell.nichd.nih.gov/. The published Seurat object was processed in R (v 4.4.0) using Seurat (v5.2.1)^69^. The R script of this analysis can be obtained at GitHub (https://github.com/MunjalLab/ccn1l1_project). Clusters of otic tissue cells from 36 hpf to 72 hpf were isolated from the original Daniocell atlas. 3000 highly variable genes were selected using the Seurat function ‘FindVariableFeatures’ with the method ‘vst’. All genes were then scaled to have a mean expression of 0 and a variance of 1 across all selected cells. Clustering was performed using the Louvain algorithm^70^ implemented in FindClusters with a resolution of 0.6. The canal-forming cell cluster was identified based on expression of *vcanb*, a known marker for this cell type, supported by previous literature^11,15^. The top 10 differentially expressed genes were identified by the ‘FindAllMarkers’ function in Seurat.

### Comparative analysis of Ccn1l1 and Ccn1 amino acid sequences

For the analysis of the functional domains of zebrafish Ccn1l1, amino acid sequences were retrieved from the NCBI database. Functional domains were predicted using the SMART database^71,72^, and the significance (E-value) of each predicted domain was recorded. To compare domain conservation across species, multiple sequence alignments were performed using Clustal Omega^73^, and sequence similarity and percent identity were assessed. Clustal alignment outputs were used to visualize conserved residues, and the pairwise identity matrix was used to quantify percent identity between species. Amino acid sequences of each functional domain predicted by SMART are listed in Supplementary Table 2.

### Analysis of the TEAD-binding domain in 5’ upstream region of *ccn1l1*

For this analysis, genomic DNA was extracted from EK wild-type embryos at 2 dpf, and a 7-kb fragment of the 5′ upstream region of *ccn1l1* was amplified by PCR. The sequence of the PCR product was determined by Sanger sequencing and analyzed for the presence of the TEAD-binding motif (GGAATG/CATTCC)^24,28–30^. The following primers were used for PCR: forward, 5′-TATGTCAAACACACAAGCAGTGG-3′; reverse, 5′-GGTGTTATCCAAAAAATTGTTAAGAGGC-3′.

### Quantification and statistical analysis

The majority of image analysis was performed in Fiji^74^, unless otherwise noted.

#### 1) Bud length measurement

For quantification, images were acquired by confocal live imaging of *Tg(actb2:membrane-neongreen-neongreen*) embryos, with z-stacks collected at 0.5 or 1 μm intervals. 2D images capturing the middle section of the bud were used to measure the bud length of embryos. Bud lengths were measured manually using the “straight-line” tool in Fiji. Where budding had not started or in perturbations where budding was affected, bud lengths correspond to average cell height at the anterior or posterior region. For time-lapse analysis, the same buds were tracked every 30 min, and the bud length was measured at each time point. For the anterior bud length of *gpr126^st49/st49^* mutants or the anterior pillar of WT embryos (Fig. 5), bud length was measured by Imaris software. Using the “spot” tool, spots were placed along the centerline of the bud, and the total length was calculated by connecting these spots.

#### 2) Cell and nuclear aspect ratio

The cell aspect ratio of budding cells was measured using the “Straight-Line” tool in Fiji. Z-stack images of the OV in *Tg(actb2:membrane-neongreen-neongreen*) embryos were acquired by confocal live imaging at 0.5 μm intervals. 2D images capturing the middle section of budding cells were used to measure cell height and width, as shown in Extended Fig. 1a. For time-lapse analysis, the same cells were tracked every 30 min, and the cell aspect ratio was measured at each time point. To measure the 3D nuclear aspect ratio (Fig. 4g), embryos were injected with H2B-EGFP mRNA, followed by confocal live imaging at 50 hpf. Z-stack images of anterior buds were acquired at 0.5 μm intervals. The 3D nuclear aspect ratio was measured using Imaris. Using the “Surface” tool, masks delineating the H2B-EGFP signal were generated at 1 μm intervals in the 3D images and subsequently merged to reconstruct an ellipsoidal 3D nuclear model, as shown in Extended Fig. 4e. The lengths of the long and short axes of the ellipsoid were quantified, and the nuclear aspect ratio was calculated. For time-lapse analysis of the 2D nuclear aspect ratio (Fig. 4i’ and Extended Fig. 4f), Z-stack images of the OV in *TgKI(Yap1-mScarlet); Tg(actb2:H2B-EGFP)* embryos were acquired by confocal live imaging at 1 μm intervals. 2D images capturing the middle section of cell nuclei were used to measure nuclear height and width, and the 2D nuclear aspect ratio of individual cells was tracked at 1 hour intervals

#### 3) Counting cell numbers in a bud

Cell counting was performed using the Mastodon plugin (https://github.com/mastodon-sc/mastodon) in Fiji, which enables 3D image analysis. 3D z-stack images of the otic vesicle in *Tg(actb2:membrane-neongreen-neongreen*) embryos were acquired by confocal live imaging at 0.5 or 1 μm slice intervals. Using the Big Data Viewer in the Mastodon plugin, all cells in a bud were manually marked in 3D and counted. For time-lapse analysis, the same buds were tracked every 30 min, and the cell number was counted at each time point. For counting the *ccn1l1-expressing* cells, Z-stack images of OV in embryos stained with HCR-FISH probe against *ccn1l1* and Hoechst 33342 were used. The number of *ccn1l1*-expressing cells was quantified counted by Imaris “spot “ tool.

#### 4) Bud-ECM volume measurement

The bud-ECM region was defined as shown in Extended Fig. 1a. 3D z-stack images of the otic vesicle in *Tg(actb2:membrane-neongreen-neongreen*) embryos were acquired by confocal live imaging at 1 μm slice intervals. Bud-ECM volume was measured using Imaris software. Using the “surface” tool, masks delineating the bud-ECM area were generated at 1 μm slice intervals in the 3D images and subsequently merged to reconstruct a 3D surface filling the bud-ECM region, and the volume was quantified. For time-lapse analysis, the same buds were tracked every 30 min, and the volume of bud-ECM region was measured at each time point.

#### 5) Intensity analysis of multiplex *in situ* HCR (HCR-FISH)

Measurement of the fluorescence intensity of the HCR probes was performed as previously described^11^. In brief, using Fiji, maximum intensity z-projections covering the entire anterior or posterior bud were created. To measure the mean intensity of HCR probes, the ROI was manually drawn along the outer cells of the bud with a “segmented line” tool. To create a histogram plot of fluorescence intensity, regions of interest (ROIs) were manually drawn using the “segmented line” tool (10 pixels wide) along the anterior budding region, spanning 55-65 μm, as shown in Fig. 2. An additional 20 μm on each side of the non-budding region flanking the budding cells was also included. The “plot profile” tool was then used to obtain an average intensity trace along the line. Fluorescence intensities were divided into 120 and 40 bins for the budding and shoulder regions, respectively, expressed as normalized distances, and subsequently normalized again to the overall maximum value. The Fire heatmap was used to represent the fluorescence intensities of individual probes. The heatmap color scale was adjusted individually for each experiment.

#### 6) Intensity analysis of the Yap1 nuclear-to-cytoplasm ratio

For measurement of the Yap1 nuclear-to-cytoplasmic ratio in immunostained samples, nuclear and cytoplasmic regions in a 2D slice at the middle section of each cell were manually segmented based on Hoechst and phalloidin staining, respectively, using Fiji software, and the ratio of mean nuclear to mean cytoplasmic signal intensities was calculated. For measurement using the *TgKI(Yap1-mScarlet)* line, nuclei in a 2D slice at the middle section of each cell were manually segmented based on the H2B-EGFP signal, and the mean nuclear Yap1 signal intensity was quantified. The mean cytoplasmic Yap1 signal intensity was obtained on the same 2D slice using the “oval” tool in Fiji by placing four circular ROIs in the cytoplasm and averaging their mean intensities. The Yap1 nuclear-to-cytoplasmic ratio was then calculated as described above. For time-lapse analysis of Yap1 nucleae-to-cytoplasmic ratio, posterior buds of *TgKI(Yap1-mSacrlet); Tg(actb2:H2B-EGFP)* embryos were used. The Yap1 nuclear-to-cytoplasmic ratio of individual cells was tracked at 1 hour intervals as described above. To generate color-coded images based on the Yap1 nuclear-to-cytoplasmic ratio, we referred to previously published methods^75^. Briefly, Z-stack images of the posterior bud from *TgKI (Yap1-mScarlet); Tg(actb2:H2B-EGFP)* embryos were reconstructed from four consecutive 2D slices. Nuclear masks for individual cells in the Z-stack images were created in Fiji based on the H2B-EGFP signal. The positional information of each nuclear mask was then extracted, and the corresponding Yap1 nuclear-to-cytoplasmic ratio values, quantified from the 2D slices, were mapped onto each nuclear mask using MATLAB to generate color-coded images.

### Measurement of the number of embryos with unfused buds

Number of embryos injected with control MO or *yap1* MO with visible unfused bud morphologies by the total number of embryos measured at 72 hpf. The “unfused” phenotype includes embryos with failure of fusion of all buds and those with fusion failure of only the posterior and ventral buds.

### Statistical analysis

Average values were calculated from ‘n,’ where ‘n’ represents the number of buds, bud-ECM, and cells (as labeled in each figure). Plots show individual data points, mean values, and error bars (standard deviation of the mean, as labeled). Before performing parametric statistical tests, the Kolmogorov-Smirnov and Shapiro-Wilk tests were used to test if the dataset in each experiment was from a normal distribution. To compare the difference between the two groups, unpaired or paired two-tailed Student’s t-tests were performed. For multiple comparisons, one-way ANOVA with Tukey’s test for normally distributed data and Kruskal-Wallis test with Dunn’s test for non-normally distributed data were performed. P-values less than 0.05 were considered to be significant. All statistical tests and plots were made in GraphPad Prism 10.

## Acknowledgments

We are thankful to Nadia Eliora, Vishank Jain-Sharma, and other members of the Principles of Tissue Morphogenesis (Munjal) Lab for their feedback and support. We thank Brigid Hogan and Stefano Di Talia for their critical comments on the manuscript. We thank Kenneth Poss for providing the *hsp70l: RFP-DN Yap1* construct. We thank members of the Poss and Bagnat labs for valuable technical input during this project. We thank Sean Megason and Ian Swinburne for their guidance and support during AM’s postdoctoral training. We thank Kazunori Ando for technical support and advice to YM. We thank Toru Kawanishi and Sean Megason for sharing the Yap immunostaining protocol. We thank Tony Tsai’s lab for sharing the *Yap (4xGTIIC:EGFP)* reporter, which was not included in the final manuscript. We thank Nanami Sato for illustrations. We thank Ziqi Lu for assistance in developing the code for generating time-lapse movies of the *TgKI(Yap1-mScarlet)* line. We thank the Duke Zebrafish Core Facility of the Duke School of Medicine for their support. This work was funded by the National Institute of Child Health and Human Development (R00HD098918 and DP2115157 to A.M.) and the National Institute of Arthritis and Musculoskeletal and Skin Diseases (R01AR083346 to M.B). YM is supported by JSPS Overseas Research Fellowship, Duke Regeneration Centre Career Advancement Award, and Uehara Memorial Foundation Overseas Research Fellowship.

## Author contributions

A.M. conceived and initiated the project. Y.M., with input from A.M., subsequently developed the project direction. Y.M. designed, performed, and analyzed most of the experiments with the assistance of A.M., A.B.B., and K.L.H. P.R. developed the theoretical model with the supervision of E.H. Y.M. generated the *Tg(hsp70l: RFP-DN-Yap1)* line, *Tg(hsp70l: DN-PKA-GFP)* line, *Tg(actb2: H2B-EGFP)* line, and all constructs for the project, except those required for the *TgKI(Yap-mScarlet)* line. D.S.L. and M.B. generated the *TgKI(Yap-mScarlet)* line and the corresponding construct. J.W. analyzed the publicly available transcriptomics dataset. Y.M. and P.R. prepared figures with the input from E.H. and A.M. Y.M. and A.M. interpreted the data and wrote the original manuscript with input from P.R. and E.H. All authors reviewed and edited the manuscript. A.M. acquired funding for the project. A.M. supervised and administered the project.

