## Supplementary Information for "A self-limiting mechanotransduction feedback loop ensures robust organ formation"

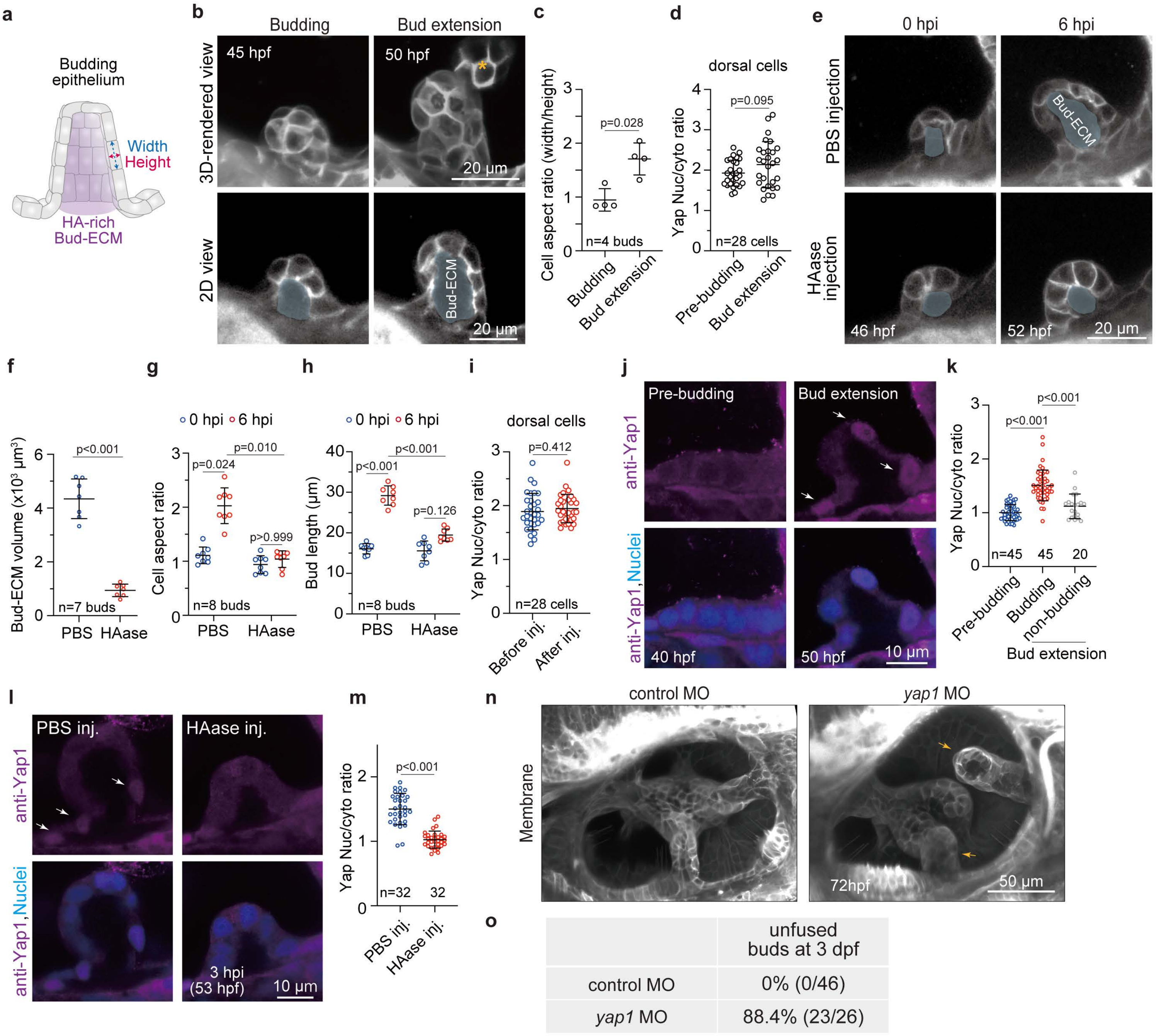


**Extended Figure 1**

**Extended Figure 1: ECM swelling-mediated Yap activation in budding cells, related to Figure 1**

(a) Illustration of budding epithelium and bud-ECM region filled with hyaluronan.

(b) 3D-rendered and 2D section of the anterior bud imaged at budding (45 hpf) and bud extension (50 hpf) stages using *Tg(actb2:membrane-neongreen-neongreen)* embryos. Yellow asterisk indicates the opposing anterior-lateral bud. Regions marked in light blue indicate the bud-ECM region. Scale bar: 20 μm. Representative images from two independent experiments are shown.

(c) Quantification of cell aspect ratio in budding cells in the anterior bud at budding (45 hpf) and bud extension (50 hpf) stages. Cell height and width in a budding cell are defined as shown in Extended Fig. 1a. Data are mean ± SD. *n* denotes the number of buds from individual embryos measured per condition from two independent experiments. *P*-value as labeled (unpaired two-tailed Student's t-test).

(d) Quantification of the Yap nuclear-to-cytoplasmic ratio in the dorsal epithelial cells at pre-budding (40 hpf) or bud extension (50 hpf) stages. Data are mean ± SD. *n* denotes the number of cells measured per condition from nine individual embryos of two independent experiments. *P*-value as labeled (unpaired two-tailed Student's t-test).

(e) 2D sections of the anterior bud in *Tg(actb2:membrane-neongreen-neongreen)* embryos imaged at 0 and 6 hpi of PBS and HAase. 0 hpi and 6 hpi images were acquired from the same embryo. Scale bar: 20 μm. Representative images from two independent experiments are shown.

(f) Quantification of bud-ECM volume of PBS or HAase-treated anterior buds at 6 hpi (52 hpf). Data are mean ± SD. *n* denotes the number of buds measured per condition from two independent experiments. *P*-value as labeled (unpaired two-tailed Student's t-test).

(g) Quantification of cell aspect ratio of budding cells in anterior buds before (46 hpf) and at 6 hpi (52 hpf) of PBS and HAase. Data are mean ± SD. *n* denotes the number of buds from individual embryos measured per condition from two independent experiments. *P*-values as labeled (Kruskal-Wallis test with Dunn's test).

(h) Quantification of bud length of anterior buds at 0 hpi (46 hpf) and at 6 hpi (52 hpf) of PBS and HAase. Data are mean ± SD. *n* denotes the number of buds from individual embryos measured per condition from two independent experiments. *P*-values as labeled (one-way ANOVA with Tukey's test).

(i) Quantification of the Yap nuclear-to-cytoplasmic ratio in the dorsal epithelial cells before (50 hpf) and after (51.5 hpf) injection of HAase. Data are mean ± SD. *n* denotes the number of cells measured per condition from nine individual embryos of two independent experiments. *P*-value as labeled (Mann-Whitney test).

(j-k) Immunostaining of Yap1 in budding cells. (j) 2D sections of anterior buds showing Yap1 and nuclear staining using anti-Yap1 antibody and Hoechst stain respectively, at selected time points. Scale bar: 10 μm. Arrows indicate Yap1 translocated to the nucleus. Representative images from two independent experiments are shown. (k) Quantification of the Yap nuclear-to-cytoplasmic ratio in the epithelial cells at the canal genesis zone at selected time points. Data are mean ± SD. *n* denotes the number of cells measured per condition from two independent experiments. *P*-values as labeled (Kruskal-Wallis test with Dunn's test).

(l-m) Effect of HAase on nuclear Yap translocation. (l) 2D sections of anterior buds showing Yap1 and nuclear staining using anti-Yap1 antibody, and Hoechst stain, respectively, at 3 hpi (53 hpf) of PBS or HAase. Scale bar: 10 μm. Arrows indicate Yap1 translocated to the nucleus. Representative images from two independent experiments are shown. (m) Quantification of the Yap nuclear-to-cytoplasmic ratio in the budding cells in the anterior buds at 3 hpi of PBS or HAase. Data are mean ± SD. *n* denotes the number of cells measured per condition from two independent experiments. *P*-value as labeled (Mann-Whitney test).

(n) 3D-rendered OVs from *Tg(actb2:membrane-neongreen-neongreen)* embryos, injected with control or *yap1* MO, imaged at 72 hpf. Yellow arrows indicate unfused posterior and ventral buds. Scale bar: 50 μm. Representative images from three independent experiments are shown.

(o) Table shows the percentage of control or *yap1* MO-injected embryos with unfused buds in OVs at 72 hpf.

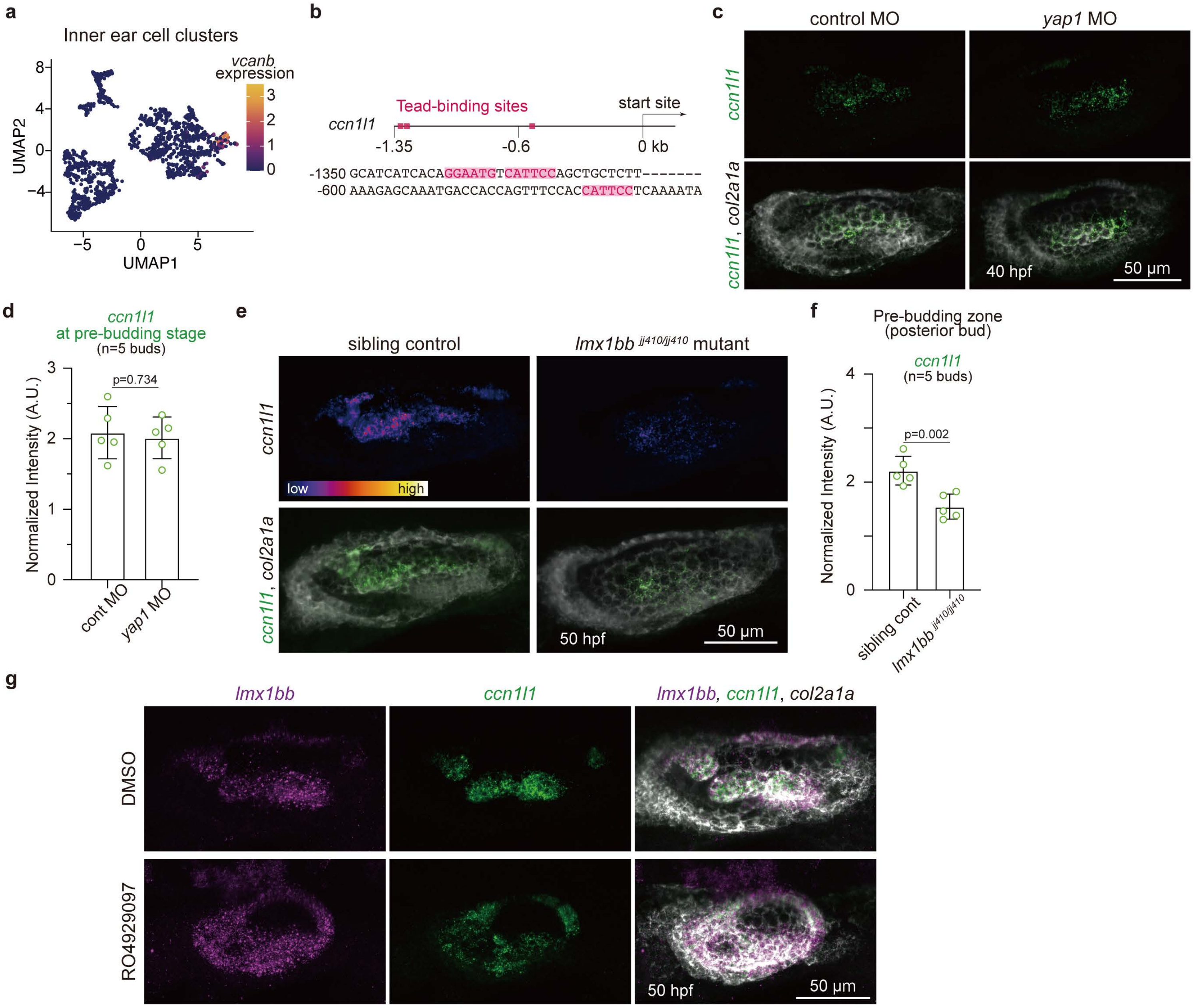


**Extended Figure 2**

**Extended Figure 2: Lmx1bb regulates *ccn1l1* expression at the canal-genesis zone, related to Figure 2**

(a) UMAP of single-cell RNA of *vcanb*-expressing cell cluster from the DanioCell dataset. Heatmap color of each dot represents *vcanb* expression levels.

(b) Schematic image and sequence of the TEAD binding motifs in the genomic sequence upstream of the *ccn1l1* locus.

(c-d) Effect of *yap1* knockdown on the onset of *ccn1l1* expression at the pre-budding stage. (c) 3D-rendered OVs, injected with control or *yap1* MO, imaged at 40 hpf stained with multiplex *in situ* probes against *ccn1l1* (green) and *col2a1a* (white). Scale bar: 50 µm. Representative images from two independent experiments are shown. (d) Quantification of probe fluorescence intensity of *ccn1l1* probe in the anterior budding region at the pre-budding stage (40 hpf). Data are mean ± SD. *n* denotes the number of budding regions from individual embryos measured from two independent experiments. *P*-value as labeled (unpaired two-tailed Student's t-test).

(e-f) Effect of *lmx1bb* mutation on *ccn1l1* expression level. (e) 3D-rendered OVs of *lmx1bb* $^{jj410/jj410}$ or sibling control embryos at 50 hpf stained with multiplex *in situ* probes against *ccn1l1* (green) and *col2a1a* (white). Heatmap represents fluorescence intensity with the same contrast for the *ccn1l1* probe of each. Scale bar: 50 µm. Representative images from three independent experiments are shown. (f) Quantification of probe fluorescence intensity of *ccn1l1* probe in the posterior pre-budding region. Data are mean ± SD. *n* denotes the number of budding regions from individual embryos measured from two independent experiments. *P*-value as labeled (unpaired two-tailed Student's t-test).

(g) Maximum intensity projections of OVs treated with Notch receptor inhibitor RO4929097 at 50 hpf stained with multiplex *in situ* probes against *lmx1bb* (magenta), *ccn1l1* (green), and *col2a1a* (white). Scale bar: 50 µm. Representative images from three independent experiments are shown.

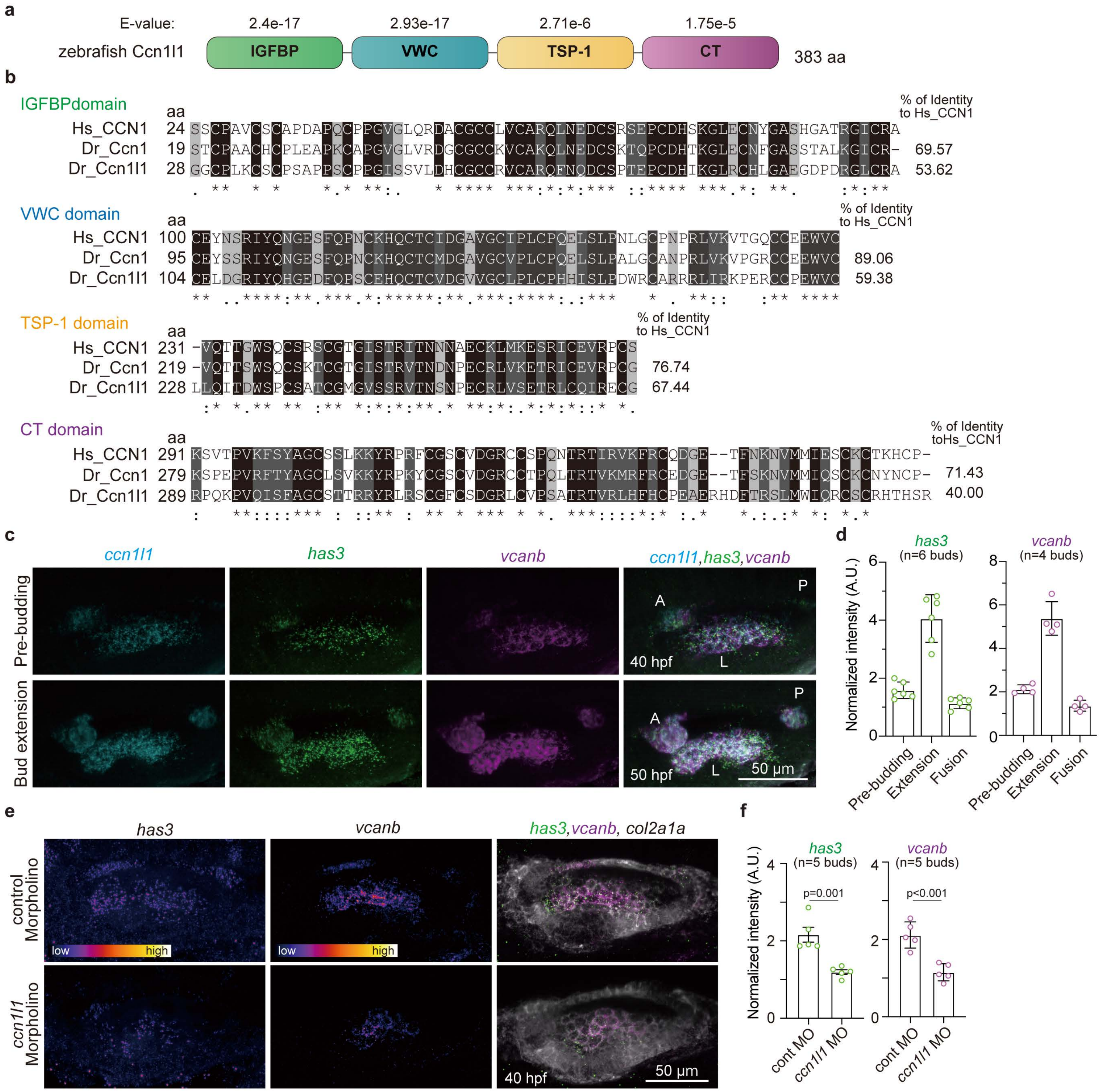


**Extended Figure 3**

**Extended Figure 3: Ccn1l1 induces *has3* and *vcanb* expression in budding cells, related to Figure 3**

(a) E-values of predicted functional domains of zebrafish Ccn1l1 obtained with SMART.

(b) Multiple sequence alignment of amino acids of each functional domain from zebrafish (*Danio rerio*) Ccn1l1 (Dr_Ccn1l1), zebrafish Ccn1 (Dr_Ccn1), and human (*Homo sapiens*) CCN1 (Hs_CCN1) performed with Clustal Omega. Black (*), dark grey (:), and grey (.) indicate fully conserved residues, strong similarity of residues, and weak similarity of residues, respectively, across species. The percentage sequence identity of Dr_Ccn1 and Dr_Ccn1l1 relative to Hs_CCN1 is shown.

(c) 3D-rendered OVs at pre-budding (40 hpf) or bud extension (50 hpf) stages stained with multiplex *in situ* probes against *ccn1l1* (cyan), *has3* (green), and *vcanb* (magenta). A, anterior bud; L, lateral bud; P, posterior bud. Scale bar: 50 μm. Representative images from three independent experiments are shown.

(d) Quantification of probe fluorescence intensities of *has3 and vcanb* probes measured at the pre-budding stage (anterior budding region), extension stage (anterior bud region), or fusion stage (anterior pillar region). Data are mean ± SD. *n* denotes the number of buds from individual embryos measured from two independent experiments.

(e-f) Effect of *ccn1l1* knockdown on *has3* and *vcanb* expression at the pre-budding stage. (e) 3D-rendered OVs injected with control or *ccn1l1* MO at 40 hpf stained with multiplex *in situ* probes against *has3* (green), *vcanb* (magenta), and *col2a1a* (white). Heatmaps represent fluorescence intensity with the same contrast for each probe of embryos. Scale bar: 50 μm. Representative images from two independent experiments are shown. (f) Quantification of probe fluorescence intensities of *has3* and *vcanb* probes in the anterior pre-budding region. Data are mean ± SD. *n* denotes the number of buds from individual embryos measured from two independent experiments. *P*-values as labeled (unpaired two-tailed Student's t-test).

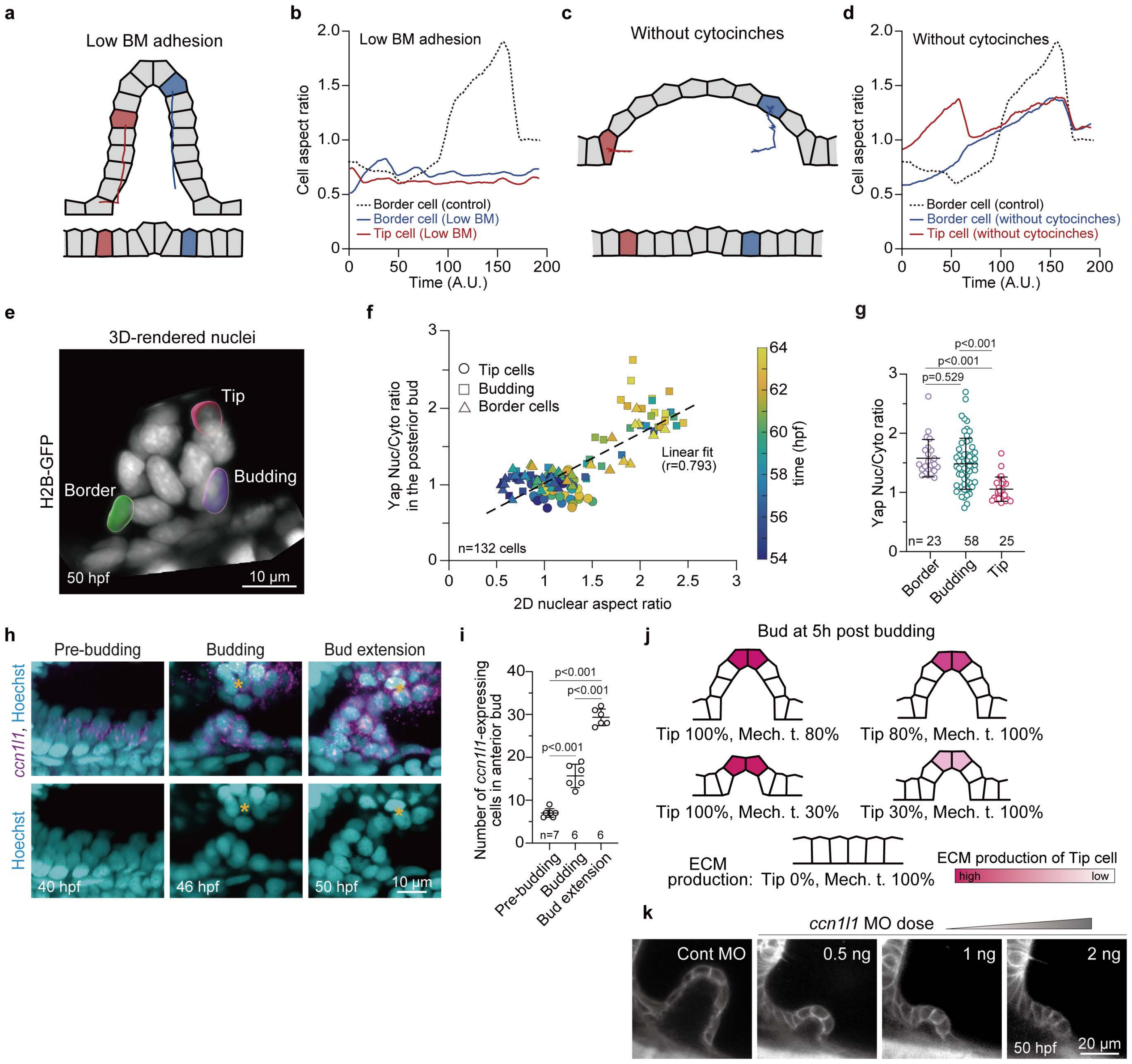


**Extended Figure 4**

**Extended Figure 4: Mechanical patterning of budding cells, related to Figure 4**

(a) Simulation snapshot of a bud with low adhesion to the basement membrane obtained with the 2D vertex model, showing the trajectory of two cells during bud extension. Upper and lower panels show the bud shape after and before budding, respectively.

(b) Simulated aspect ratio of a tip cell and a recruited cell during bud extension, in the case of low adhesion to the basement membrane. The dashed black line indicates the trajectory of the aspect ratio of a recruited cell in the control case.

(c) Simulation snapshot of a bud in the absence of cytocinches obtained with the 2D vertex model, showing the trajectory of two cells during bud extension. Cells appear to be pushed away from the bud center, rather than flowing towards it. Upper and lower panels show the bud shape after and before budding, respectively.

(d) Simulated Aspect ratio of a tip cell and a recruited cell during bud extension obtained with the 2D vertex model, in the absence of cytocinches. The dashed black line indicates the trajectory of the aspect ratio of a recruited cell in the control case.

(e) 3D-rendered nuclei of an anterior bud from *Tg(actb2:H2B-EGFP)* embryos at 50 hpf. Representative nuclear masks for tip, budding, and border cells are shown. Scale bar: 10 μm. Representative images are shown from two independent experiments.

(f) Correlation of Yap nuclear-to-cytoplasmic ratio and 2D nuclear aspect ratio in tip, budding, and border cells of a posterior bud across individual time points obtained from time-lapse imaging of *TgKI(Yap1-mScarlet)*; *Tg(actb2:H2B-EGFP)* embryos. Heatmap represents developmental time. *n* denotes the number of cells from four buds from three independent experiments. Dashed line indicates linear fit (Pearson correlation coefficient: $r = 0.793$).

(g) Yap nuclear-to-cytoplasmic ratios in border, budding and tip cells within the anterior bud at 50 hpf. Data are mean ± SD. *n* denotes the number of cells per condition from two independent experiments. *P*-values as labeled (one-way ANOVA with Tukey's test).

(h) 3D-rendered anterior buds at selected time points stained with *in situ* probe against *ccn1l1* (magenta) and Hoechst stain (cyan). Yellow asterisks indicate the opposing anterior-lateral bud. Scale bar: 10 μm. Representative images from three independent experiments are shown.

(i) Quantification of number of *ccn1l1* expressing cells in anterior buds at selected time points. Number of *ccn1l1* expressing cells was counted using embryos stained with *in situ* probe against *ccn1l1* and Hoechst stain. *n* denotes the number of buds of individual embryos measured from two independent experiments. *P*-values as labeled (one-way ANOVA with Tukey's test).

(j) Simulation snapshots of bud morphology at 5 hours post-budding for different ECM production rate parameter values. The red heatmap shows the ECM production rate in tip cells. Mech. t. indicates the percentage of ECM production driven by mechanotransduction contribution from recruited cells.

(k) 2D sections of anterior buds from *Tg(actb2:membrane-neongreen-neongreen)* embryos injected with control or *ccn1l1* MO at the indicated concentrations, imaged at 50 hpf. Scale bar: 20 µm. Representative images from two independent experiments are shown.

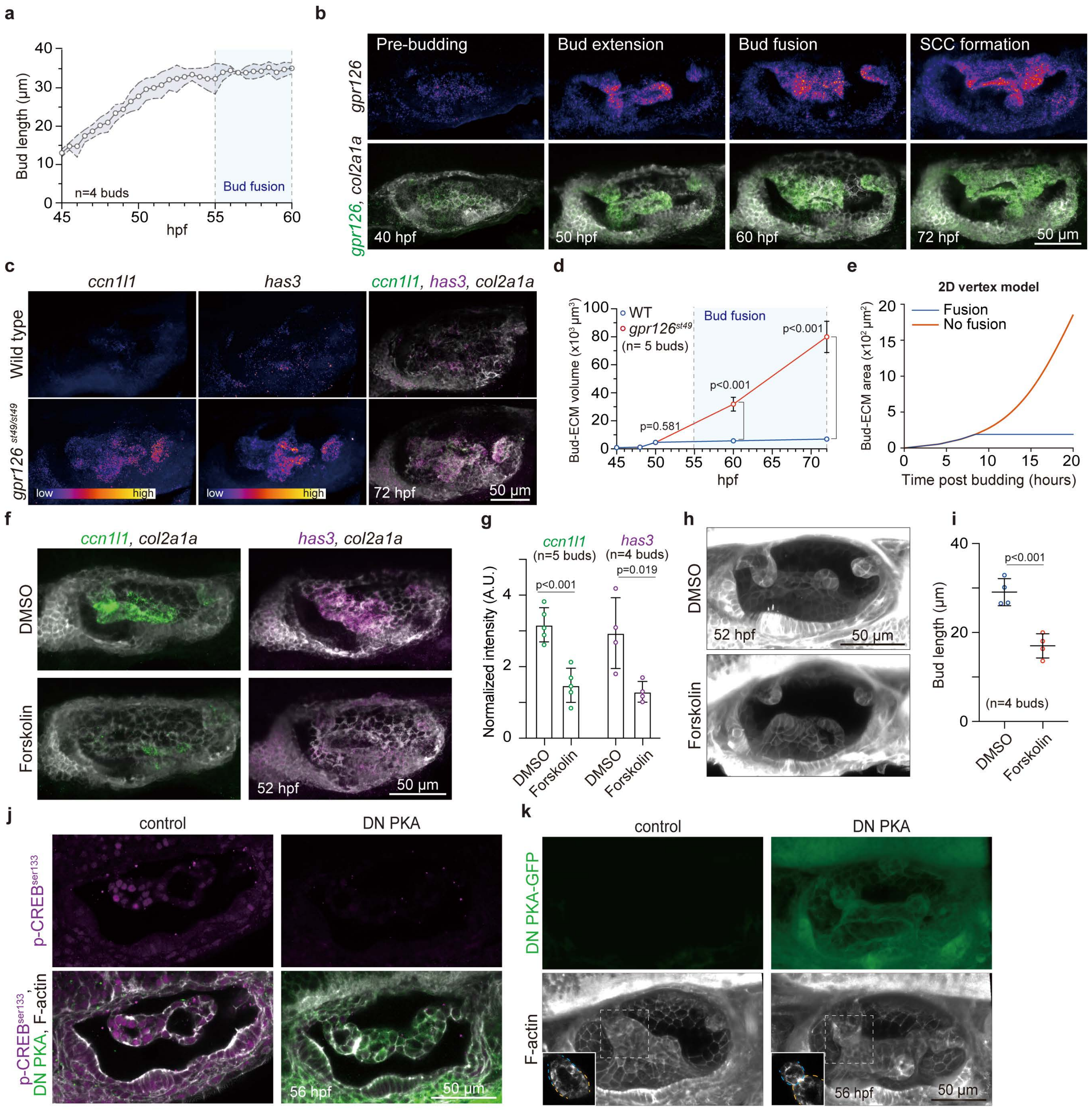


Extended Figure 5

**Extended Figure 5: Gpr126-cAMP-PKA-CREB signaling terminates bud extension, related to Figure 5**

(a) Quantification of anterior bud length across individual time points obtained from time-lapse imaging of *Tg(actb2:membrane-neongreen-neongreen)* embryos. Data are mean ± SD. Dashed lines indicate the range of SD at each time point. *n* denotes the number of buds from individual embryos measured from three independent experiments.

(b) 3D-rendered OVs at selected time points stained with multiplex *in situ* probes against *gpr126* (green) and *col2a1a* (white). Heatmap represents fluorescence intensity with the same contrast for the *gpr126* probe of each embryo. Scale bar: 50 μm. Representative images from two independent experiments are shown.

(c) 3D-rendered OVs of wild-type and *gpr126* $^{st49/st49}$ mutant embryos at 72 hpf stained with multiplex *in situ* probes against *ccn1l1* (green), *has3* (magenta), and *col2a1a* (white). Heatmaps represent fluorescence intensity with the same contrast for each probe of wild-type or *gpr126* $^{st49/st49}$ mutant embryos. Scale bar: 50 μm. Representative images from three independent experiments are shown.

(d) Quantification of bud-ECM volume in anterior buds across individual time points obtained from time-lapse imaging of wild-type or *gpr126* $^{st49/st49}$ embryos, both *Tg(actb2:membrane-neongreen-neongreen*). Data are mean ± SD. *n* denotes the number of buds from individual embryos measured from three independent experiments. *P*-values as labeled (unpaired two-tailed Student's t-test)

(e) Simulated bud ECM area (equivalent of bud ECM volume in our 2D simulation) as a function of time, with (fusion: blue line) or in the absence (no fusion: red line) of the terminating event of bud fusion. Tip and budding cell productions are set to 100%.

(f-g) Effect of forskolin treatment on *ccn1l1* and *has3* expression. (f) 3D-rendered OVs of embryos treated with forskolin at 52 hpf stained with multiplex *in situ* probes against the combination of *ccn1l1* (green) and *col2a1a* (white) or *has3* (magenta) and *col2a1a* (white). Scale bar: 50 μm. Representative images from two independent experiments are shown. (g) Quantification of probe fluorescence intensities of the *ccn1l1* and *has3* probes in the anterior bud. Data are mean ± SD. *n* denotes the number of buds from individual embryos measured from two independent experiments. *P*-values as labeled (unpaired two-tailed Student's t-test).

(h-i) Effect of forskolin treatment on bud growth. (h) 3D-rendered OVs of *Tg(actb2:membrane-neongreen-neongreen)* embryos, treated with forskolin, imaged at 52 hpf. Scale bar: 50 μm. Representative images from two independent experiments are shown. (i) Quantification of anterior bud length in embryos treated with forskolin at 52 hpf. Data are mean ± SD. *n* denotes the number of buds from individual embryos measured per condition from two independent experiments. *P*-value as labeled (unpaired two-tailed Student's t-test).

(j) 2D sections of OVs showing phospho-CREB, DN PKA-GFP and F-actin staining using anti-phospho-CREB$^{ser133}$ antibody, anti-GFP antibody, and phalloidin, respectively, at 56 hpf in sibling control or in DN PKA-GFP-expressing embryos following heat-shock induction. Scale bar: 50 μm. Representative images from two independent experiments are shown.

(k) 3D-rendered OVs from *Tg(hsp70l: DN PKA-GFP)* and sibling control embryos, both *Tg(βActin: utrophin-mCherry)*, imaged at 56 hpf following heat-shock induction. Insets show 2D sections of the pillar or unfused buds from the regions in the white dashed boxes. Blue and yellow dashed lines indicate the anterior bud/pillar and the anterior-lateral bud/pillar, respectively. Scale bar: 50 μm. Representative images from two independent experiments are shown.

**Extended Movie 1: Mechanical epithelial cell recruitment during bud extension, related to Figure 4.**

3D-rendered time-lapse images of the anterior bud obtained from *a Tg(actb2:membrane-neongreen-neongreen)* embryo. Each cell is marked by uniquely colored dot. Images were captured every 30 min. Time stamps indicate developmental time (hours post fertilization). Scale bar: 20 µm. Fig. 4b shows a cropped frame from this movie.

**Extended Movie 2: Spatiotemporal dynamics of Yap activation in successively recruited epithelial cells during bud extension, related to Figure 4.**

Time-lapse analysis of the Yap1 nuclear-to-cytoplasmic (N/C) ratio in a posterior bud of *TgKI(Yap1-mScarlet)*; *Tg(actb2:H2B-EGFP)* embryos. Yap1 N/C ratio (left), Yap1-mScarlet (center), and H2B-EGFP (right) are shown, respectively. Z-stack images were used for Yap1-mScarlet and H2B-EGFP. Heatmap indicates Yap1 N/C ratio. Yap1 N/C ratios in tip cells (t) and border cells (b) were tracked. Arrows indicate border cells that incorporated into the posterior bud. Images were captured every 1 hour. Time stamps indicate developmental time (hours post fertilization). Scale bar: 20 µm. Fig. 4h shows cropped frames from this movie.

**Extended Movie 3: Semicircular canal morphogenesis of wild-type and *gpr126* $^{st49/st49}$ mutant embryos, related to Figure 5.**

3D-rendered time-lapse images of the OV showing SCC formation of a wild-type or a *gpr126* $^{st49/st49}$ mutant, both *Tg(actb2:membrane-neongreen-neongreen*. Images were captured every 30 min. Time stamps indicate developmental time (hours post fertilization). Scale bar: 50 µm.

### Supplementary Information – Theory note

In these notes, we detail the vertex models used to account for the bud morphogenesis, as well as analytical models to describe the role of the mechanotranduction feedback. These notes are organized as follows. In Section 1, we present the implementation of the vertex models. In Section 2, we focus on the elongation of budding cells during the recruitment process, in relation with YAP activation. In Section 3, we investigate the growth dynamics of the bud in presence of a feedback between YAP mechanotransduction and gel production.

#### 1 Methods: Vertex model

##### 1.1 Bud mechanics

As the bud is axisymmetric, we use here a 2D vertex model, representing a transversal cut across the bud in the $xy$ plane. Because the radius of the bud is much smaller than the total size of the otic vesicle, we model the later as a planar surface at $y = 0$, from which the bud is growing. The system consists in $N$ cells (corresponding to cells already recruited into the bud, and several rows of cells neighbouring the bud). The energy of a given cell $i$ is taken to be:

$$e_i = (\gamma_a \ell_a + \gamma_b \ell_b + \gamma_l \ell_l) + K(a - a_0)^2 + \mathcal{U}_{\text{adh}} + \mathcal{U}_{\text{cytocinches}}, \tag{1}$$

where $\ell_a$, $\ell_b$ and $\ell_l$ are the apical, basal and lateral lengths of the cell, respectively, and the $\gamma$'s the respective mechanical tensions from the acto-myosin cytoskeleton. The second term imposes volume conservation, with $a$ the 2D area of the cell, $a_0$ its homeostatis area and $K$ a constant energy penalty for breaking volume conservation. To account for the corner-like shapes cells go through in vivo during the recruitment process, each cell is modelled by an hexagon, with 4 vertices corresponding to contact points with adjacent cells (2 on the apical side, and 2 on the basal side), and one additional vertex on both the apical and the basal sides, to allow cells to bend. The third term corresponds to an adhesion energy, for non-budded cells still in contact with the otic vesicle:

$$\mathcal{U}_{\text{adh}} = -G \times d_{\text{contact}} e^{-y/y_{\text{contact}}}, \tag{2}$$

where $G$ is an adhesion energy per unit of contact length, $d_{\text{contact}}$ is the portion of the cell's basal length that lies below some fixed threshold $y_{\text{contact}}$, and $y$ is the average height of the cell's basal side. The exponential term allows for a smooth delamination of cells from the substrate, rather than a sharp threshold effect.

The last term in $e_i$ accounts for cytocinches, actin-rich cables that were previously determined to constrict the bud and make it anisotropic, we applied a constricting force on all vertices that are part of the bud at a given timepoint (that is, that are not in contact with the horizontal substrate):

$$\mathcal{U}_{\text{cytocinches}} = \sum_{\text{basal vertex } j} k_c x_j^2, \tag{3}$$

where the sum runs other all the basal vertices of the cell, with $x_j$ their horizontal position.

The total energy of the bud is then:

$$\mathcal{E} = K_{\text{ECM}}(A_{\text{ECM}} - A_0(t))^2 + \sum_{\text{cell } i=1}^{N} e_i, \tag{4}$$

where the first term imposes the volume of the ECM below the bud, with its equilibrium 2D area $A_0(t)$ being updated at each timestep to account for ECM production in the bud.

We then simulate the bud's mechanics through a Monte Carlo algorithm. At each timestep, we try optimizing the position and shape of each cell by randomly moving their vertices, and checking if the proposed change leads to a decrease in the total energy of the bud. In the initial state, cells on the horiizontal line are placed such that the epithelium is tension-free: each cell has a width $w = \sqrt{a \frac{\gamma_l}{\gamma_a + \gamma_b - G}}$ and height $h = a_0/w$, where $a_0$ is the homeostatic 2D area of a cell. The bud is assumed to have an axis of symmetry at its center, which in our simulation corresponds to the right-most vertices (the other half of the bud being obtained by axial symmetry). The position of the left-most vertex is kept constant throughout the simulation. All the codes were implemented in Julia.

#### 1.2 Values of parameters

Cytoskeletal tensions can be estimated by considering cellular aspect ratios. At the start of the bud extension phase, when pressure from the ECM is still low, cells are likely close to their prefered aspect ratio, given by $\epsilon_0 = w_0/h_0 = \gamma_l/(\gamma_a + \gamma_b)$, with $w_0$ and $h_0$ the width and thickness of cells in their stress-free state. We measured that their thickness was approximately $h = 9\,\mu\text{m}$ and their width $w = 5\,\mu\text{m}$, resulting in an aspect ratio $\epsilon = w/h \simeq 0.55$. We chose $\gamma_l, \gamma_a$ and $\gamma_b = 1$ to be all equal as this choice would yield $\epsilon_0 = 0.5$. In practice, experimental observations show that myosin is preferentially localized on the basal side; however, it is difficult to discriminate how much this basal myosin contributes directly to contraction, rather than simply to cell stiffness. In addition, cytocinches also contribute to contraction on the basal side. Therefore, we decided to keep $\gamma_b = \gamma_a = 1$ rather than considering more complex scenarios, and tuned basal contraction only through cytocinches.

We chose to work with length units so that the cell cross-section is $a_0 = 1$. Experimentally, the cell cross-section at the start of the bud extension phase was roughly $a_0 = 50\,\mu\text{m}^2$, so a unit length in our simulation corresponds to a distance $\sqrt{a_0} = 7\,\mu\text{m}$ in experiments.

The last two parameters, $G$ and $k_c$, representing substrate adhesion and cytocinches contractility, respectively, cannot be directly determined experimentally. Intuitively, the corresponding forces must be of the same order as the in-plane tissue tension $\gamma_a + \gamma_b$, and their precise values could be obtained by fitting the length and radius of the bud for different parameter values on experimental data. However, the 2D and highly simplified nature of our model makes such a fitting procedure rather questionable. Instead, we chose to set $G = 0.5$ and $k_c = 0.2$ in most cases, and we tested the impact of changing these values on the bud morphology.

In the following, we considered two separate cases for the dynamics of the ECM production, defined by $A_0(t)$. As a first step, we externally prescribed $A_0$ to slowly increase at a constant rate, to focus on cell recruitment alone. Then, we considered the role of the mechanotransduction feedback, and assume that the growth rate $\dot{A}_0$ was proportional to the number of recruited cells.

When testing the effects of a low adhesion, we set $G = 0.3$. In simulations without cytocinches, we set $k_c = 0.0$.

### 2 Cell elongation during the recruitment process

#### 2.1 Link between cellular aspect ratio and mechanical forces

At the lowest possible degree of description, apical, basal and lateral cytoskeletal tensions create a spring-like restoring force, causing each cell to spontaneously return to a prefered aspect ratio. For a rectangular 2D cell, this prefered value can be determined as:

$$\epsilon_0 = \frac{w_0}{h_0} = \frac{w_0^2}{a_0} = \frac{\gamma_l}{\gamma_a + \gamma_b}, \tag{5}$$

where $\epsilon_0$ is the prefered aspect ratio, $w_0$ the prefered width, $h_0$ the height of the cell, $a_0$ its 2D cross-section, and $\gamma_l$, $\gamma_a$, $\gamma_b$ are the lateral, apical and basal tensions. As a result, a deformed cell, with aspect ratio $\epsilon > \epsilon_0$ will exert a pulling force on its environment, given by:

$$\zeta = \gamma_a + \gamma_b - \gamma_l \epsilon^{-1} \simeq (\gamma_a + \gamma_b)\frac{w - w_0}{w_0}, \tag{6}$$

so that this pulling force is proportional to the relative elongation of the cell. Therefore, in what follows, we monitor the aspect ratio $\epsilon = \frac{w^2}{a}$ as a readout of mechanical forces felt by the cells. When $\epsilon > \epsilon_0$, the cell is under stretch, and this can typically result in nuclear relocalization of Yap.

#### 2.2 Cell aspect ratio during recruitment

Based on this result, we monitor the cellular aspect ratio of all cells in our simulations. First, we assume that bud ECM production is prescribed independently from cell recruitment, and we impose that $A_0 = rt$, with $r$ a constant growth rate. As cells are not rectangular, we define numerically $w$ as the average of the apical and basal sides of the cell, and compute the aspect ratio through $\epsilon = w^2/a_0$. While other definitions of the aspect ratio are possible, this one directly relates to mechanical forces as it results from the balance between apical, basal, and lateral tensions with external forces from the surrounding tissue.

Numerically, we find that cells undergo a transient stretching as they are recruited into the bud: border cells, that are partially recruited and partially still attached to the basement membrane exhibit an overshoot in their aspect ratio, and similarly for budding cells (bud cells that are adjacent to a border cell). On the other hand, tip cells showed a lower aspect ratio. Looking at trajectories of individual cells, we find that they “flow” towards the bud, and start stretching as they turn the corner. Bud adjacent cells (cells neighboring border cells, but still attached to the basement membrane) displayed a slight compression. Intuitively, we would expect that the flow of cells converging around the bud would result in an in-plane tension that decreases with distance from the bud, resulting instead to a slight stretching of bud adjacent cells. However, our 2D simulation does not model the fluidity (through in-plane rearrangements and friction with the substrate) of the epithelium around the bud. Therefore, we assume that this slight compression of bud adjacent cell stems from a small simulation artefact.

We hypothesized that the increase in aspect ratio of border cells was the result of a tug-of-war between adhesion forces, which pull the epithelium towards the basement membrane, and tugging forces from the cytocinches. To test this idea, we considered the effects of either reducing focal adhesion, or removing forces from cytocinches. The former led to over-recruitment of cells. The latter, on the other hand, fully suppressed the converging flow of cells towards the bud. Instead, the bud appeared to radially "push" the epithelium away from the center. In both cases, the aspect ratio of cells remained low and no marked difference was found between border cells and tip cells.

### 3 Role of the feedback loop

Next, we consider the impact of this mechanical patterning on the bud extension dynamics. Rather than prescribing the volume increase of the bud ECM, we now assume its rate depends on the number of recruited cells. We consider the following model:

$$\frac{\mathrm{d}A_{\mathrm{ECM}}}{\mathrm{d}t} = k(aN_{\mathrm{tip}} + bN_{\mathrm{recruited}}), \tag{7}$$

where bud ECM production is the sum of two contributions: a constant rate from tip cells, and a variable contribution from recruited cells that grows as the bud recruits more and more cells. Setting $b = 0$ allows us to mimick Yap morpholino conditions, while $b = a = 1$ would be closer to wild type conditions. Here, the quantity $k$ is a unit of ECM volume production per unit of time, setting the unit of time of our simulation. Here we choose $k = 13\,\mu\mathrm{m}^3/\mathrm{h}$, with one hour corresponding to 30000 simulation steps.

Numerically, we find that the mechanical feedback increases the bud extension rate, as well as the cell recruitment rate. The tell-sign of the feedback is the growth dynamics of the bud-ECM volume $A_{\mathrm{ECM}}$: in absence of any feedback, it grows linearly and slowly, while in its presence it grows supra-linearly. This observation is fully in line with experimental findings. In particular, in the gpr126 mutant, the bud ECM volume displays a clear non linear growth as the feedback continues unimpeded in absence of bud fusion.

To study the impact of the feedback loop on bud fusion, we define bud fusion as the moment when the bud reaches 30 µm in length. The start of our simulation corresponds roughly to a timepoint of 50 hpf. If the bud does not reach this value after 22 hours (72 hpf), we assume that the bud fails to fuse.
